# iDePP: a genetically encoded system for the inducible depletion of PI(4,5)P_2_ in *Arabidopsis thaliana*

**DOI:** 10.1101/2020.05.13.091470

**Authors:** Mehdi Doumane, Léia Colin, Alexis Lebecq, Aurélie Fangain, Joseph Bareille, Olivier Hamant, Youssef Belkhadir, Yvon Jaillais, Marie-Cécile Caillaud

## Abstract

Phosphatidylinositol 4,5-bisphosphate [PI(4,5)P_2_] is a low abundant lipid present at the plasma membrane of eukaryotic cells. Extensive studies in animal cells revealed the pleiotropic functions of PI(4,5)P_2_. In plant cells, PI(4,5)P_2_ is involved in various cellular processes including the regulation of cell polarity and tip growth, clathrin-mediated endocytosis, polar auxin transport, actin dynamics or membrane-contact sites. To date, most studies investigating the role of PI(4,5)P_2_ in plants have relied on mutants lacking enzymes responsible for PI(4,5)P_2_ synthesis and degradation. However, such genetic perturbations only allow steady-state analysis of plants undergoing their life cycle in PI(4,5)P_2_ deficient conditions and the corresponding mutants are likely to induce a range of non-causal (untargeted) effects driven by compensatory mechanisms. In addition, there are no small molecule inhibitors that are available in plants to specifically block the production of this lipid. Thus, there is currently no system to fine tune PI(4,5)P_2_ content in plant cells. Here we report a genetically encoded and inducible synthetic system, iDePP (Inducible <u>De</u>pletion of <u>P</u>I(4,5)P_2_ in <u>P</u>lants), that efficiently removes PI(4,5)P_2_ from the plasma membrane in different organs of *Arabidopsis thaliana*, including root meristem, root hair and shoot apical meristem. We show that iDePP allows the inducible depletion of PI(4,5)P_2_ in less than three hours. Using this strategy, we reveal that PI(4,5)P_2_ is critical for cortical microtubule organization. Together, we propose that iDePP is a simple and efficient genetic tool to test the importance of PI(4,5)P_2_ in given cellular or developmental responses but also to evaluate the importance of this lipid in protein localization.

**Research Organism:** *A. thaliana*

## INTRODUCTION

Phosphoinositides are “tiny lipids with giant impacts” on cell biology (Balla, 2013). These low abundant membrane lipids are derived from phosphatidylinositol. Phosphatidylinositol’s head can be phosphorylated on positions 3 and/or 4 and/or 5, leading to the synthesis of up to seven different phosphoinositide species. Detection of phosphoinositides revealed that each species is differentially enriched in the various membranes of eukaryotic cells (Platre and Jaillais, 2016), constituting a landmark code that allows specific targeting of proteins to given membranes (Bigay and Antonny, 2012; Simon et al., 2016). In particular, phosphatidylinositol 4,5-bisphosphate [PI(4,5)P_2_] is mainly present in the cytosolic leaflet of the plasma membrane in wild-type cells of *Saccharomyces cerevisiae* (Eumycota Ascomycota; Patton and Lester, 1992), human (Metazoa Mammal; Hammond et al., 2009, 2012), *Drosophila melanogaster* (Metazoa Hexapoda; El Kadhi et al., 2011), and *Arabidopsis thaliana* (Archaeplastida Angiosperm; Simon et al., 2014; Van Leeuwen et al., 2007). Pharmacological and genetically encoded systems allowing to exogenously modulate PI(4,5)P_2_ levels in animal cells led to significant advances toward the comprehensive understanding of PI(4,5)P_2_ functions (Idevall-Hagren and De Camilli, 2015). While phosphoinositide manipulation via inhibitors and/or activators of enzymes involved in phosphoinositide metabolism is easily achievable in vitro, the use of these drugs in cell cultures is often constrained by the permeability of the plasma membrane which in turn limits their activities to narrow ranges of concentrations. Thus, the use of such molecules is even less tractable on pluricellular organisms. Second, drugs often have unknown off-target effects that affect cellular processes distinct from phosphoinositide metabolism. With these limitations in mind, the pharmacological targeting of phosphoinositide kinases, phosphatases and phospholipases is nevertheless a valuable tool to study, relatively fast, the effects of phosphoinositide levels modulation. Genetically-encoded fast and reversible systems are privileged for the study of the role of the phosphoinositide in animal cell culture, however there is still a striking gap between state-of-the-art acute phosphoinositide manipulation carried out in cultured cells and those carried out in pluricellular organisms.

At the plasma membrane of animal cells, PI(4,5)P_2_ has a wide spectra of actions. It is for example a precursor for second messengers such as inositol 1,4,5-trisphosphate (IP_3_), diacylglycerol (DAG) and PI(3,4,5)P_3_ secondary messengers (Antal and Newton, 2013). PI(4,5)P_2_ can be seen as a second messenger itself, since it recruits a host of proteins to the plasma membrane in a highly controlled and regulated manner (Rajala and Anderson, 2010). It can also activate ion channels, and it regulates many cellular processes including clathrin-mediated endocytosis, exocytosis, cell polarity, membrane contact sites and actin cytoskeleton remodeling (Hammond et al., 2012; Schmid and Mettlen, 2013; Yin and Janmey, 2003). Typically, PI(4,5)P_2_ is known to promote the formation of actin filament structures— beneath the plasma membrane by inhibiting or activating distinct sets of proteins (Senju et al., 2017). Together with phosphatidylinositol 4-phosphate (PI4P), PI(4,5)P_2_ also contributes to plasma membrane surface charge (Hammond et al., 2012), a basic property of the plasma membrane, which recruits many proteins through electrostatic interaction (Platre and Jaillais, 2017).

In plant cells, less is known about PI(4,5)P_2_. Intriguingly, it seems that some functions of PI(4,5)P_2_ in animals are not conserved in plants. For example, plasma membrane surface charges relies on PI4P, phosphatidylserine (PS) and phosphatidic acid (PA), but not PI(4,5)P_2_ (Simon et al., 2016a; Platre and Jaillais, 2017). Yet, mutants in PI4P 5-kinases (PIP5K) suggest that PI(4,5)P_2_ production is essential and has critical roles in development, immunity and reproduction (Heilmann, 2016; Noack and Jaillais, 2017a). However, i) the synthesis of each phosphoinositide species is tied to each other and to other lipids, ii) PI(4,5)P_2_ has pleiotropic cellular functions and iii) PI(4,5)P_2_ interacts with many different proteins. Thus, it is likely that steady-state depletion of PI(4,5)P_2_ triggers a range of indirect effects involving several types cellular and developmental compensatory mechanisms. It is therefore essential to manipulate the levels of this lipid in a rapid manner if we want to untangle direct and indirect functions of PI(4,5)P_2_. However, there are limited tools available for researcher to modulate PI(4,5)P_2_ levels in plant cells and organs in an inducible way. To date, most of the available tools allow to increase the amount of PI(4,5)P_2_. This includes U-73122, a pan-phospholipases C (PLC) inhibitor (Barbosa et al., 2016; Lee et al., 2019; Van Leeuwen et al., 2007). However, because phosphoinositide-specific PLCs (PI-PLC) use both PI(4,5)P_2_ and PI4P as substrates in plants, such treatment is likely to also impact the levels of both lipids (Delage et al., 2012). Another method is the exogenous treatment to membrane permeable PI(4,5)P_2_ analogues (Ozaki et al., 2000). However, the stability of these lipids is questionable given that phospholipids tend to be degraded into DAG upon exogenous treatment (Grabski et al., 1993). Finally, inducible expression of PIP5Ks efficiently increase the level of PI(4,5)P_2_ in plants (Gujas et al., 2017a; Ischebeck et al., 2013). Yet, to date, a strategy that allows to specifically and inducibly deplete PI(4,5)P_2_ in plants remains to be implemented. Indeed, long treatment with the PI4K inhibitor phenylarsine oxide (PAO) reduces both PI(4,5)P_2_ and PI4P levels (Platre et al., 2018; Simon et al., 2016) and there is no PIP5K inhibitor available (Kusano et al., 2008; Mei et al., 2012; Menzel et al., 2019; Stanislas et al., 2018).

Here, we built a system for the inducible depletion PI(4,5)P_2_ in plants, that we coined iDePP (<u>I</u>nducible <u>Dep</u>letion of <u>P</u>I(4,5)P_2_ in <u>P</u>lants). iDePP is a dexamethasone inducible genetically encoded system that allows fast and specific PI(4,5)P_2_ depletion in an entire organism, the model plant *Arabidopsis*. This system triggers the expression of a plasma membrane anchored PI(4,5)P_2_ 5-phosphatase domain from the *Drosophila melanogaster* OCRL upon dexamethasone treatment. Using iDePP, we detected a strong and specific PI(4,5)P_2_ depletion starting as soon as 90 min after induction. We show that the iDePP system is suitable to study the function of PI(4,5)P_2_ in both above and below-ground plant in *Arabidopsis*. Finally, we use iDePP to provide evidence that the appropriate cortical-microtubule orientation requires PI(4,5)P_2_ in the *Arabidopsis* root epidermis.

## RESULTS

### dOCRL phosphatase domain acts as a PI(4,5)P2 phosphatase *in vitro*

In cultured mammalian cells, an inducible system for the depletion PI(4,5)P_2_ was previously design (Idevall-Hagren et al., 2012). They used a truncation (amino acids 234 to 539) of human OCRL (OCRL1) corresponding to its PI(4,5)P_2_ 5-phosphatase domain. We were concerned that this domain, that underwent natural selection in an endotherm organism with set-point temperature of 37°C, would be less active in *Arabidopsis thaliana*, a plant of temperate climate which we usually grow at 21°C. Therefore, we decided to use the phosphatase domain of dOCRL, *Drosophila melanogaster* homolog, as *Drosophila* are ectotherm and live at ∼ 20°C. Alignment of OCRL1_234-539_ and dOCRL allowed us to identify dOCRL_168-509_ as the homolog region of OCRL1 generally used in optogenetic systems (Figure 1A). We first assessed dOCRL activity, which has not been reported so far, even though *in vivo* data suggest a conserved PI(4,5)P_2_ 5-phosphatase activity (Carim et al., 2019; Ben El Kadhi et al., 2011; Del Signore et al., 2017). We ordered a synthetic gene corresponding to dOCRL_168-509_ and codon optimized for expression in *Arabidopsis*. We cloned a recombinant dOCRL protein (His-SUMO-dOCRL_168-509_-His) tagged with two 6-histidine tags and a SUMO tag, making it suitable for expression in *Escherichia coli* and purification. We purified His-SUMO-dOCRL_168-509_-His using a cobalt resin-based affinity chromatography and protein concentration (Figure 1B). We subjected the purified protein fractions C1 and C2 to Malachite Green Phosphatase assays, that detects phosphate released by dephosphorylations, in presence of short chain (diC:8) water-soluble phosphoinositides (Maehama et al., 2000). Note that we only assessed dOCRL_168-509_ activity toward phosphoinositides reported in plant cells (Platre and Jaillais, 2016; Simon et al., 2014). We observed that dOCRL_168-509_ displayed a strong and specific phosphatase activity toward PI(4,5)P_2_ (Figure 1C), demonstrating that dOCRL activity is conserved between Mammals and *Drosophila*. Therefore, dOCRL_168-509_ is a domain suitable for specific PI(4,5)P_2_ depletion at room temperature (∼ 20°C).

**Figure 1.**
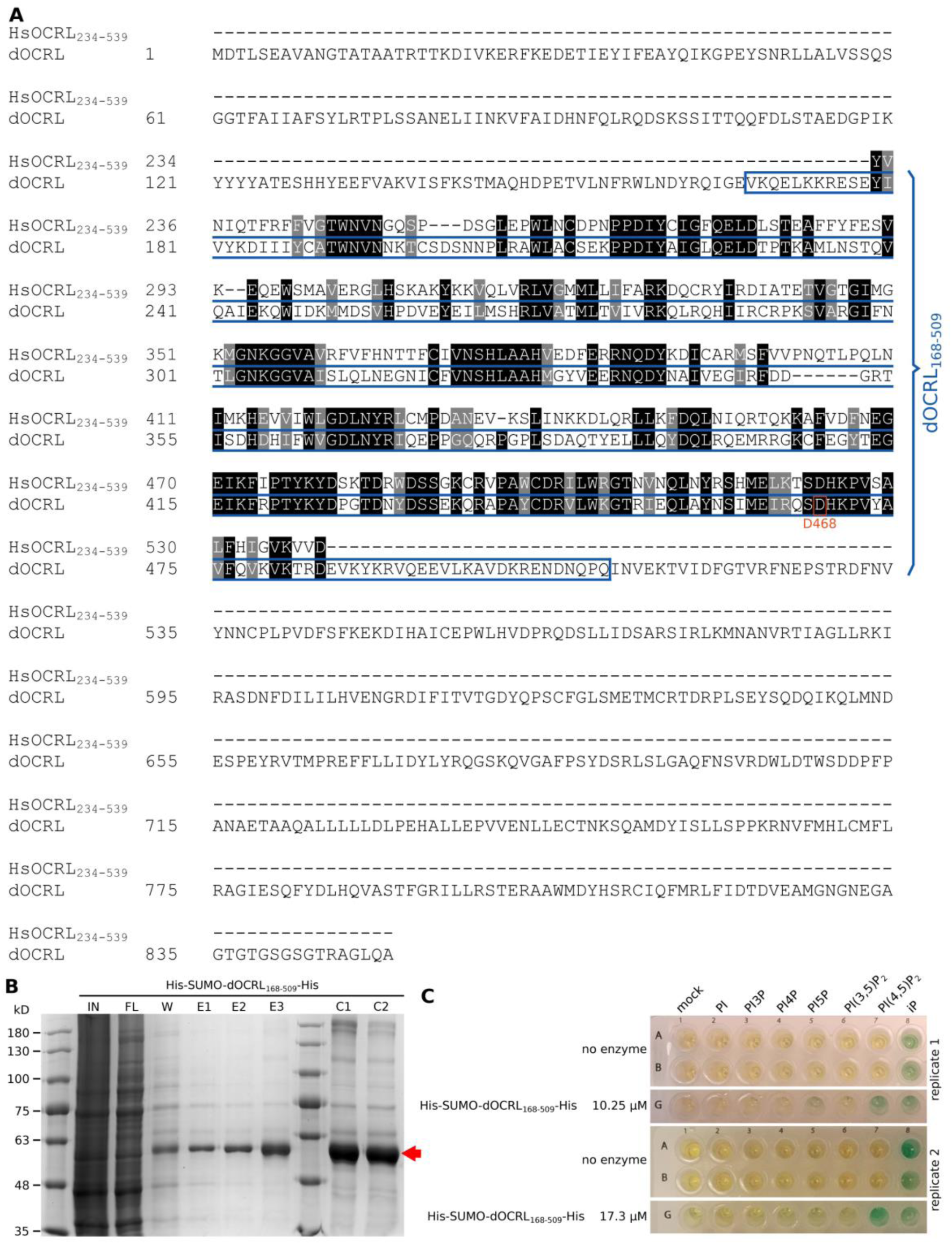
*Drosophila* dOCRL displays a PI(4,5)P_2_ phosphatase activity in vitro. (A) Alignment of (i) *Homo sapiens* OCRL (HsOCRL or OCRL1) truncation (HsOCRL_234-539_) successfully used in optogenetic systems in mammalian cells (Idevall-Hagren et al., 2012), and (ii) of *Drosophila melanogaster* dOCRL. We cloned dOCRL_168-509_ truncation, framed in blue, that is homolog to HsOCRL_234-539_. Aspartate 468 (D468), framed in red, is the amino acid we mutated to obtain inactive dOCRL_168-509_ (see Figure 2 and 3). (B) Coomassie blue staining monitoring purification of recombinant dOCRL phosphatase domain (His-SUMO-dOCRL_168-509_-His) from *Escherichia coli*. IN: input, lysate of bacteria induced for the expression of His-SUMO-dOCRL_168-509_-His; FL: flow through; W: wash; E1 to E3: elution fraction 1 to 3; C1 and C2: concentrated fractions from pulled E1, E2 and E3. C1 and C2 were obtained separately on different days. His-SUMO-dOCRL_168-509_-His expected molecular weight is 54.8 kD. (C) Malachite green phosphatase assay on His-SUMO-dOCRL_169-509_-His using short chain water-soluble phosphoinositides. Replicate 1 correspond to C1 fraction. Replicate 2 to C2 fractions. Mock: no phosphoinositide added.

### Inducible expression and subcellular targeting of dOCRL

After validating *in vitro* the enzymatic activity of the catalytic domain of dOCRL_168-509_, we next addressed whether this enzyme could efficiently hydrolyze PI(4,5)P_2_ in plant cell when targeted to the plasma membrane. We fused the codon optimized phosphatase domain (here after named dOCRL) to a Myristoylation And Palmitoylation (MAP) sequence, which is responsible for membrane targeting (Martinière et al., 2012) and to mCHERRY (mCH) fluorescent protein to monitor protein expression and subcellular localization of MAP-mCHERRY-dOCRL (Figure 2A). In order to observe short term effects of PI(4,5)P_2_ depletion, we took advantage of dexamethasone (dex) - GVG system that allows efficient inducible expression of proteins in *Arabidopsis* (Aoyama and Chua, 1997; Marquès-Bueno et al., 2011; Moreno-Romero et al., 2008). We next performed site-directed D468G mutation in dOCRL catalytic domain, similarly to D523G mutation already described in human OCRL1 that abolishes its phosphatase activity (Idevall-Hagren et al., 2012). We could therefore use MAP-mCH-dOCRL_168-159_^D468G^ (hereafter named MAP-mCH-dOCRL^dead^) as negative control, together with a MAP-3xmCH recombinant protein (Figure 2A).

**Figure 2.**
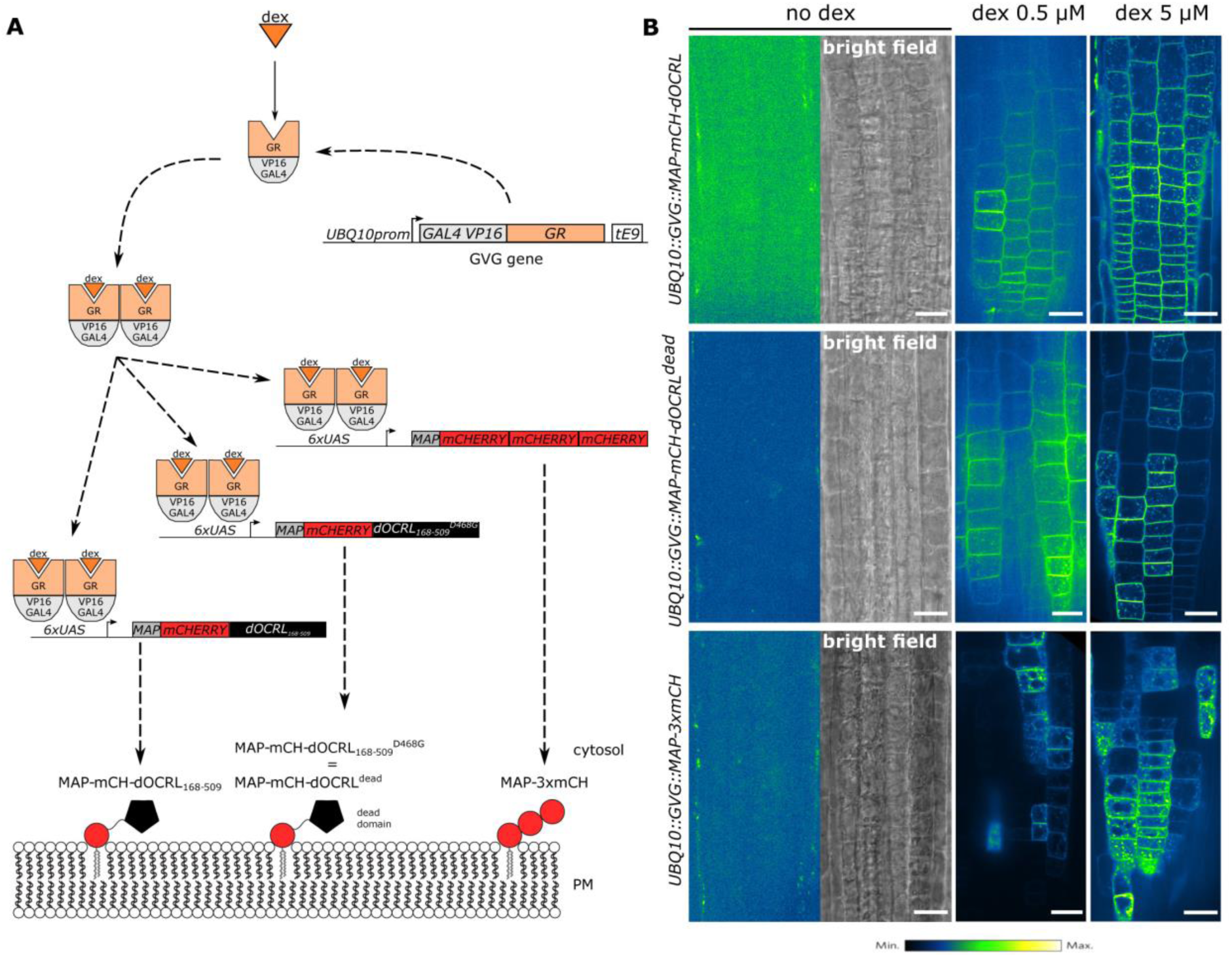
Successful inducible expression and subcellular targeting of MAP-mCH-dOCRL and negative controls. (A) Inducible system used in this study. *UBQ10* promoter is responsible for mild/strong and ubiquitous expression of *GVG* (*GAL4 VP16 GR*) synthetic gene. GR domain binds dexamethasone (dex) and subsequently induces GVG homodimerization and nuclear import. There, GAL4 domain binds *UAS* DNA elements and VP16 strongly activates the expression of downstream gene: *MAP-mCH-dOCRL*_*168-509*_, or *MAP-mCH-dOCRL*_*168-509*_^*D468G*^ (*MAP-mCH-dOCRL*^*dead*^), or *MAP-3xmCH*. MAP is myristoylation and palmytoylation sequence, responsible for plasma membrane targeting. mCH corresponds to monomeric CHERRY fluorescent protein. dOCRL^D468G^_168-509_ is an inactive phosphatase domain (later on called dOCRL^dead^). 3xmCH correspond to three fused mCHERRY. (B) For all three inducible constructs we monitored mCH fluorescence without dex treatment or after a 16h-treatment with either 0.5 µM or 5 µM dex. Signal intensity is color-coded (green fire blue scale). Scale bars: 20 µm.

We next addressed the timing of expression and the localization of each construct in *Arabidopsis* lines stably transformed with *UBQ10pro:GVG:MAP-mCH-dOCRL*, or *UBQ10pro:GVG:MAP-mCH-dOCRL*^*dead*^, or *UBQ10pro:GVG:MAP-3xmCH*. In root meristematic epidermal cells, without dex treatment, none of the recombinant-proteins were detected using confocal microscopy (Figure 2B). A 16 h treatment with 0.5 µM dex led to the detection of mCH fluorescence, indicating that induction of the genetic construct had occurred (Figure 2B). However, we observed a mosaic induction and a significant number of roots had no cells expressing the fluorescent reporter to detectable levels. To overcome these issues, we optimized the treatment to a 16 h induction with 5 µM dex. Using this set up, we robustly observed red fluorescence using confocal microscopy where MAP-mCH-dOCRL, MAP-mCH-dOCRL^dead^ and MAP-3xmCH (Figure 2B). Each of the synthetic protein was efficiently targeted to membranes, including the plasma membrane, where PI(4,5)P_2_ accumulates, and intracellular compartments (Figure 2B). Therefore, a 16h 5µM dex treatment is sufficient for an effective expression of dOCRL in *Arabidopsis* stable transgenic lines.

### Inducible expression of MAP-mCH-dOCRL specifically and efficiently depletes PI(4,5)P_2_ in stable *Arabidopsis* lines

Next, we tested if the targeting of dOCRL at the plasma membrane was actively depleting PI(4,5)P_2_. For this, we monitored the localization of genetically encoded fluorescent biosensors, made of a lipid-binding domain (LBD) fused to a fluorescent protein such as mCITRINE (mCIT), which allow monitoring lipid subcellular localization and levels, using confocal microscopy (Colin and Jaillais, 2020). We previously reported that upon phosphatidylinositol 4-phosphate (PI4P) and phosphatidylserine (PS) depletion from the plasma membrane, PI4P and PS biosensors are partially or totally released from the plasma membrane, and are found in the cytosol, their default localization in the absence of a cognate lipid (Platre et al., 2018; Simon et al., 2016). Consistent with this idea, we observed a partial release of the mCIT-2xPH^PLC^ PI(4,5)P_2_ biosensor from the plasma membrane into the cytosol upon long-term treatment with PAO, a known inhibitor of PI4P and PI(4,5)P_2_ synthesis (Simon et al., 2016). We thus predicted that the PI(4,5)P_2_ biosensors would behave similarly upon MAP-mCH-dOCRL expression.

To test this, we generated transgenic lines stably expressing various mCITRINE-tagged anionic lipid sensors with the *MAP-mCH-dOCRL* line. First, we studied the impact of MAP-mCH-dOCRL induction on the localization of the high affinity mCIT-2xPH^PLC^ PI(4,5)P_2_ biosensors (Simon et al., 2014), in *Arabidopsis* meristematic cells. In the absence of dex, MAP-mCH-dOCRL was not expressed, and only background signal was observed (Figure 3A, left panel), while mCIT-2xPH^PLC^ mainly localized to the plasma membrane and was only weakly found in the cytosol (Figure 3A, right panel). 16h of dex treatment resulted in the expression of MAP-mCH-dOCRL and the full solubilization of mCIT-2xPH^PLC^ (Figure 3B and G, dissociation indexes of ∼ 2, Supplementary file 1-3), indicating an efficient depletion of PI(4,5)P_2_ from the plasma membrane. Next, we crossed the mCIT-2xPH^PLC^ reporter line with the MAP-mCH-dOCRL^dead^ and MAP-3xmCH control lines and tested the impact of the induction of these controls on the plasma membrane targeting of the PI(4,5)P_2_ reporter. In each case, the localization of the mCIT-2xPH^PLC^ biosensor was not affected and remained at the plasma membrane either in the absence or presence of dex induction (dissociation indexes of ∼ 1 and Figure 3 C, 3G Supplementary file 1-3).

**Figure 3.**
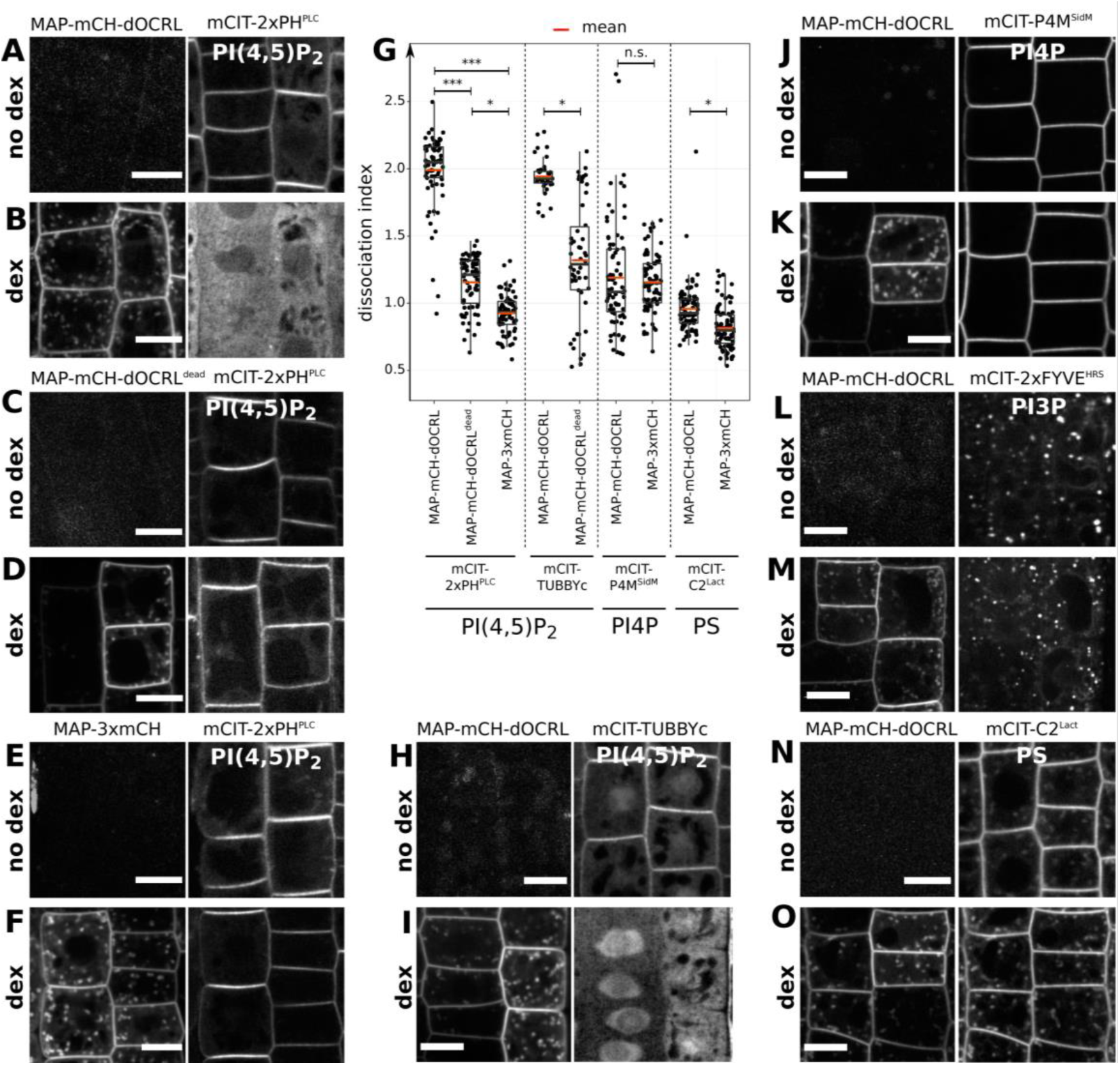
Fast, inducible and specific depletion of PI(4,5)P_2_ in plant cells using iDePP. (A) Representative images of the fluorescent signal corresponding to MAP-mCH-dOCRL (left panel) and the PI(4,5)P_2_ sensor mCIT-2xPH^PLC^ (right panel) in the same root cells, without dex treatment. (B) Representative images of the fluorescent signal corresponding to MAP-mCH-dOCRL (left panel) and the PI(4,5)P_2_ sensor mCIT-2xPH^PLC^ (right panel) in the same root cells, after 16h of dex treatment. (C) Representative images of the fluorescent signal corresponding to the negative control MAP-mCH-dOCRL^dead^ (left panel) and the PI(4,5)P_2_ sensor mCIT-2xPH^PLC^ (right panel) in the same root cells, without dex treatment. (D) Representative images of the fluorescent signal corresponding to MAP-mCH-dOCRL^dead^ (left panel) and the PI(4,5)P_2_ sensor mCIT-2xPH^PLC^ (right panel) in the same root cells, after 16h of dex treatment. (E) Representative images of the fluorescent signal corresponding to MAP-3xmCH (left panel) and the PI(4,5)P_2_ sensor mCIT-2xPH^PLC^ (right panel) in the same root cells, without dex treatment. (F) Representative images of the fluorescent signal corresponding to MAP-3xmCH (left panel) and the PI(4,5)P_2_ sensor mCIT-2xPH^PLC^ (right panel) in the same root cells, after 16h of dex treatment. Scale bars: 10 µm. (G) Dissociation indexes of lipid biosensors upon expression of MAP-mCH-dOCRL or negative controls. Dissociation index is the ratio of (i) plasma membrane to cytosol fluorescence ratio without dex treatment, to (ii) plasma membrane to cytosol fluorescence ratio after dex treatment. Statistical analysis with LMER (Type II Wald χ^2^ test) and post-hoc tests (See Supplemental file 1 for details). ***: p-value < 0.0005; *: p-value < 0.05; n. s.: p-value ≥ 0.05. See Supplemental file 3 for details. (H) Representative images of the fluorescent signal corresponding to MAP-mCH-dOCRL (left panel) and the PI(4,5)P_2_ sensor mCIT-TUBBY-C (right panel) in the same root cells, without dex treatment. (I) Representative images of the fluorescent signal corresponding to MAP-mCH-dOCRL (left panel) and the PI(4,5)P_2_ sensor mCIT-TUBBY-C (right panel) in the same root cells, after 16h of dex treatment. (J) Representative images of the fluorescent signal corresponding to MAP-mCH-dOCRL (left panel) and the PI4P sensor mCIT-P4M^SidM^ (right panel) in the same root cells, without dex treatment. (K) Representative images of the fluorescent signal corresponding to MAP-mCH-dOCRL (left panel) and the PI4P sensor mCIT-P4M^SidM^ (right panel) in the same root cells, after 16h of dex treatment. (L) Representative images of the fluorescent signal corresponding to MAP-mCH-dOCRL (left panel) and the PI3P biosensor mCIT-FYVE^HRS^ (right panel) in the same root cells, without dex treatment. (M) Representative images of the fluorescent signal corresponding to MAP-mCH-dOCRL (left panel) and the PI3P biosensor mCIT-FYVE^HRS^ (right panel) in the same root cells, after 16h of dex treatment. (N) Representative images of the fluorescent signal corresponding to MAP-mCH-dOCRL (left panel) and the phosphatidylserine (PS) biosensor mCIT-C2^Lact^ (right panel) in the same root cells, without dex treatment. (O) Representative images of the fluorescent signal corresponding to MAP-mCH-dOCRL (left panel) and the phosphatidylserine (PS) biosensor mCIT-C2^Lact^ (right panel) in the same root cells, after 16h of dex treatment. Scale bars: 10 µm.

To confirm the results obtained with mCIT-2xPH^PLC^, we crossed the *MAP-mCH-dOCRL* expressing line with the *mCIT-TUBBY-C* line, which expresses an independent PI(4,5)P_2_ biosensor (Simon et al., 2014). As previously described, mCIT-TUBBY-C localized to the plasma membrane, the cytosol and the nucleoplasm in the absence of dex induction (Figure 3H). However, similar to the mCIT-2xPH^PLC^ biosensor, mCIT-TUBBY-C was removed from the plasma membrane upon 16 hours induction of *MAP-mCH-dOCRL* expression (Figure 3G and I, Supplementary file 1-3).

We next tested whether the inducible expression of *MAP-mCH-dOCRL* would affect the subcellular accumulation of other anionic lipids. To this end, we first crossed the *MAP-mCH-dOCRL* expressing line with mCIT-P4M^SidM^, which reports on plasma membrane PI4P (Simon et al., 2016). We observed that mCIT-P4M^SidM^ had the same localization in the absence or presence of *MAP-mCH-dOCRL* expression (dissociation index ∼ 1 and Figure 3G, J and K, Supplementary file 1-3). Next, we crossed *MAP-mCH-dOCRL* with the mCIT-2xFYVE^HRS^ reporter line, which express a PI3P biosensors targeted to late endosomes and the tonoplast (Simon et al., 2014). We confirmed that in the absence of dex, mCIT-2xFYVE^HRS^ was localized in intracellular compartments and weakly at the tonoplast (Figure 3L) and that we obtained a similar localization pattern upon dex treatment and induction of MAP-mCH-dOCRL (Figure 3M).

Finally, we crossed *MAP-mCH-dOCRL* with a reporter line expressing the phosphatidylserine biosensor mCIT-C2^Lact^ (Platre et al., 2018). mCIT-C2^Lact^ accumulates at the plasma membrane and all along the endocytic pathways (Platre et al., 2018), and this pattern appeared to be conserved in both the absence or presence of dex (Figure 3 N and O, Supplementary file 1-3). However, the dissociation index of mCIT-C2^Lact^ upon expression of *MAP-mCH-dOCRL* was slightly but significantly higher than upon expression of the 3xmCH control (dissociation index= of 0.957 versus 0.815, p-value = 0.043 and Figure 3G; Supplementary file 1-3), suggesting a possible impact of the depletion of PI(4,5)P_2_ on the homeostasis of phosphatidylserine.

Overall, our results show that MAP-mCH-dOCRL efficiently, specifically, and directly hydrolyzes the PI(4,5)P_2_ pool at the plasma membrane of plant cells. We named this synthetic genetic system iDePP for Inducible Depletion of PI(4,5)P_2_ in Plants.

### iDePP quickly depletes PI(4,5)P_2_ *in vivo*

To challenge the ability of MAP-mCH-dOCRL to rapidly deplete PI(4,5)P_2_, we performed a dual color time-course analysis during dex induction, monitoring mCIT-2xPH^PLC^ PI(4,5)P_2_ biosensor subcellular localization and MAP-mCH-dOCRL appearance at the root tip in epidermal cells. We used an automated root tracking set-up we previously developed (Doumane et al., 2017) and imaged root epidermal cells every five minutes for 6 h 30 min. 15min after dex treatment (first time-point), mCIT-2xPH^PLC^ labelled the plasma membrane while no signal relative to MAP-mCH-dOCRL fluorescence was observed (Figure 4A). After 90 min, a fraction of mCIT-PH^PLC^ started to be partially solubilized in the cytosol of some cells, indicating that PI(4,5)P_2_ had started to be depleted from the targeted membrane. (Figure 4A, arrows). After 180 min, mCIT-2xPH^PLC^ was cytosolic in most cells, indicating near complete PI(4,5)P_2_ depletion Over the next few hours, mCIT-2xPH^PLC^ retained a cytosolic localization while MAP-mCH-dOCRL fluorescence increased. We measured dissociation indexes of mCIT-2xPH^PLC^ over-time during this time-lapse. We observed that dissociation index increased strongly between 90 and 180 min of dex exposure, before plateauing at ∼ 1.8 (Figure 4B). At 180 min, dissociation indexes had increased in all cells, and at 330 min it had reached its maximum in all cells. Altogether, these results showed that iDePP is rapid and efficient to deplete in an inducible manner PI(4,5)P_2_ at the plasma membrane of *Arabidopsis* root cells.

**Figure 4.**
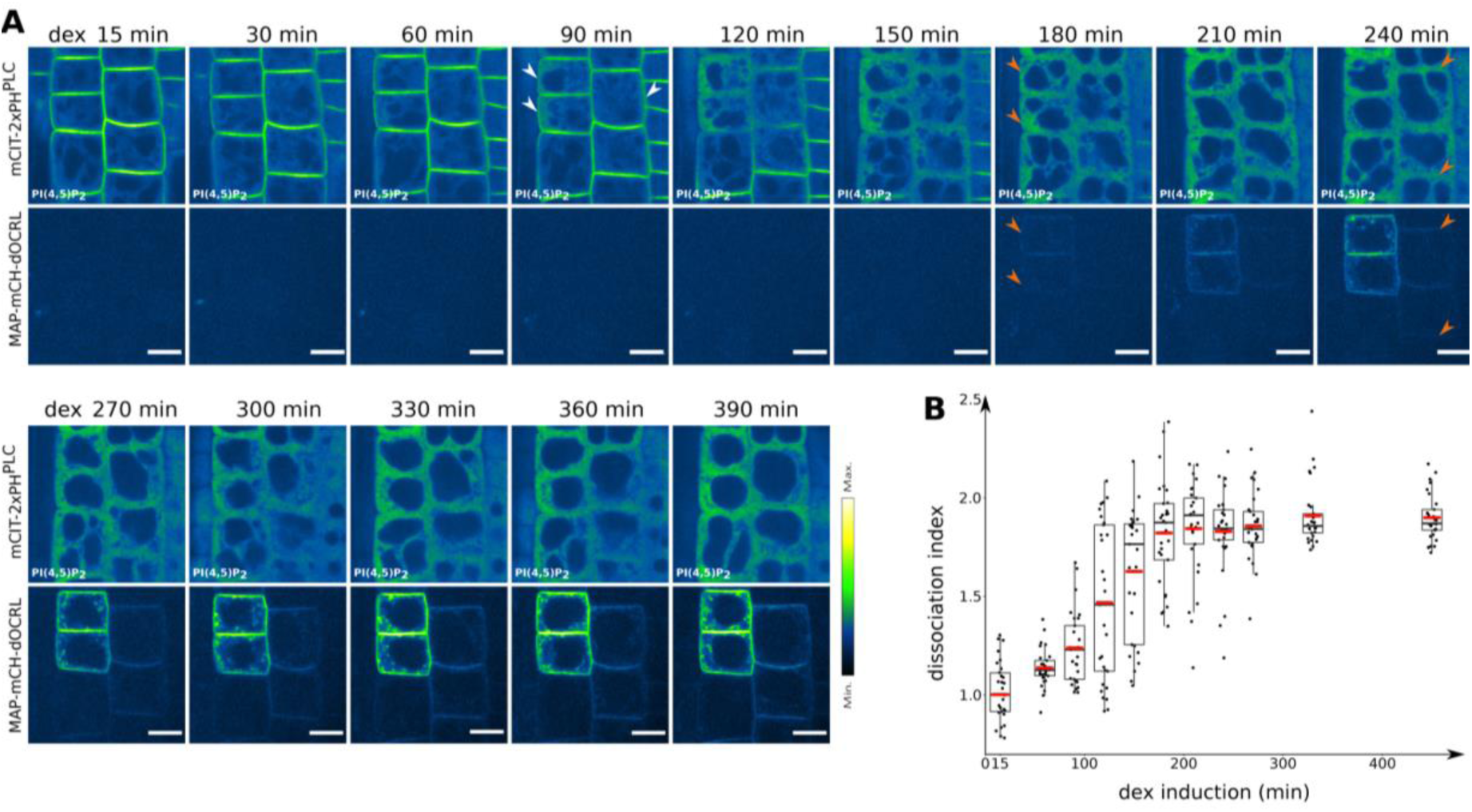
Time-lapse analysis of the depletion of PI(4,5)P_2_ in plant cells using iDePP. Dual color time-course analysis during dex induction, monitoring mCIT-2xPH^PLC^ PI(4,5)P_2_ biosensor subcellular localization and MAP-mCH-dOCRL appearance at the root tip in epidermal cells every five minutes for 6h30. 15min after the beginning of the induction, mCIT-2xPH^PLC^ labelled the plasma membrane while no signal relative to MAP-mCH-dOCRL fluorescence was observed. After 90 min, a fraction of mCIT-PH^PLC^ started to be partially solubilized in the cytosol of some cells, indicating that PI(4,5)P_2_ had started to be depleted from the targeted membrane (arrows). After 180 min, mCIT-2xPH^PLC^ was cytosolic in most cells, indicating near complete PI(4,5)P_2_ depletion (arrows). (B) Dissociation indexes of mCIT-2xPH^PLC^ over-time. Dissociation index is the ratio of (i) plasma membrane to cytosol fluorescence ratio without dex treatment, to (ii) plasma membrane to cytosol fluorescence ratio after dex treatment.

### iDePP depletes PI(4,5)P_2_ in various organs

We next determined if our system was suitable for PI(4,5)P_2_ depletion in other cell types than meristematic root epidermal cells. We therefore tested iDePP in root hairs, cotyledons and shoot apical meristems for which the PI(4,5)P_2_ subcellular and tissue patterning is well described (Noack and Jaillais, 2017; Simon et al., 2014; Stanislas et al., 2018, Colin and Jaillais 2019). Upon dex treatment, all three MAP-mCH-dOCRL, MAP-mCH-dOCRL^dead^ and MAP-3xmCH were expressed in root hairs, and localized to the plasma membrane and intracellular compartments (Figure 5A, B). Root hairs accumulate PI(4,5)P_2_ on the plasma membrane at their tip (Denninger et al., 2019; Kusano et al., 2008; Stenzel et al., 2008; Van Leeuwen et al., 2007; Vincent et al., 2005) and expression of MAP-3xmCH did not affected plasma membrane localization of mCIT-2xPH^PLC^, which remained enriched at the tip of the cell (Figure 5A, B). Conversely, expression of MAP-mCH-dOCRL solubilized mCIT-2xPH^PLC^ from the plasma membrane into the cytoplasm of root hair cells, indicating efficient PI(4,5)P_2_ depletion (Figure 5A, B).

**Figure 5.**
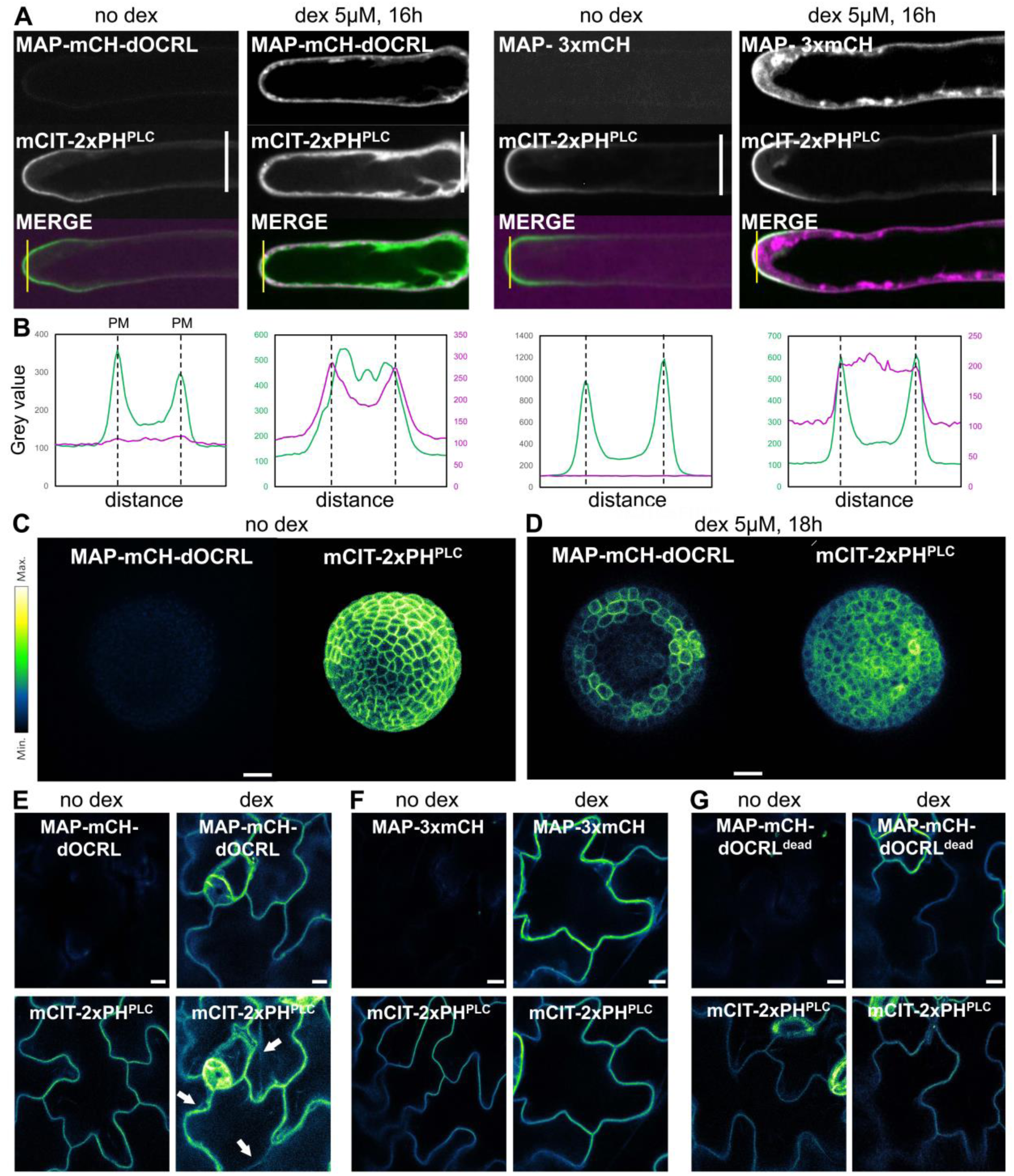
iDePP enables efficient depletion of PI(4,5)P_2_ in various tissues and organs. (A) PI(4,5)P_2_ depletion in root hairs. Subcellular localization of mCIT-2xPH^PLC^, a PI(4,5)P_2_ biosensor, and either MAP-mCH-dOCRL or MAP-3xmCH without and with 5 µM dexamethasone treatment for 16h. (B) Signal intensity for mCIT-2xPH^PLC^ (green) and MAP-mCH-dOCRL or MAP-3xmCH (magenta) with and without 5 µM dexamethasone treatment for 16h, along the yellow line drawn on panel (A). The peaks correspond to plasma membrane (PM) positions, as indicated. Note that for the condition with dex, the signal for mCH channel was weaker that with mCIT and therefore signal intensity was plotted with a different scale, on a secondary axe on the right. (C) and (D) Subcellular localization of MAP-mCH-dOCRL (left panel) and mCIT-2xPH^PLC^ (right panel) in a naked shoot meristem from an NPA-treated seedling, as viewed from the top. Absence of treatment (no dex, C) or 16 h with 5 µM dexamethasone treatment (dex, D). Scale bar, 20 μm (E to G) Subcellular localization of mCIT-2xPH^PLC^ and either MAP-mCH-dOCRL (E), or MAP-3xmCH (F), MAP-mCH-dOCRL^dead^(G), in cotyledon epidermal cells. Fluorescence intensity is color-coded (green fire blue scale), cytoplasmic strands are indicated by the arrows. Scale bars: 10 µm.

We next investigated if the iDePP system was suitable to deplete PI(4,5)P_2_ in the aerial part of the plant. For this, we first analyzed iDePP’s effects in the shoot apical meristem. Upon dex treatment, MAP-mCH-dOCRL was mainly induced in the boundary region of the shoot apical meristem and had a weak or no expression in the central zone (Supplemental file 4). This result was consistent with our previous observations, since *MAP-mCH-dOCRL* is induced in the *UBQ10* promoter expression domain, which is low at the center of the shoot apical meristem in our conditions (Stanislas et al., 2018). We observed a strong correlation between *MAP-mCH-dOCRL* expression and its impact on mCIT-2xPH^PLC^ localization (Supplemental file 4). Indeed, in the center of the shoot meristem, *MAP-mCH-dOCRL* was not expressed and mCIT-2xPH^PLC^ labelling clearly accumulated at the cell periphery, likely representing the plasma membrane (Supplemental file 4). By contrast, in the boundary zone, where *MAP-mCH-dOCRL* was expressed, mCIT-2xPH^PLC^ was not localized sharply at the cell edge and instead filled the entire volume of the cell except the nucleus (Supplemental file 4). This suggested that, similar to root epidermis, expression of *MAP-mCH-dOCRL* in the shoot epidermis displaced mCIT-2xPH^PLC^ from the plasma membrane into the cytosol, and thus efficiently erased PI(4,5)P_2_ from the plasma membrane.

Because *UBQ10*-driven *MAP-mCH-dOCRL* induction was not uniform at the shoot apical meristem, we next decided to test the iDePP system on N-1-Naphthylphthalamic Acid (NPA) grown shoot meristems. NPA is a polar auxin transport inhibitor, which induces naked shoot apical meristem without any organ (Sassi et al., 2012, Okada et al., 1991). In this condition, the *UBQ10* promoter is more uniformly expressed throughout the meristem, including the central and peripheral zones, as exemplified by the homogenous expression of *mCIT-2xPH*^*PLC*^ (Figure 5C). In the absence of dex induction, mCIT-2xPH^PLC^ was localized at the plasma membrane throughout the meristem, including both the central and peripheral zones (Figure 5C). By contrast, upon induction of *MAP-mCH-dOCRL* expression, mCIT-2xPH^PLC^ was no longer sharply localizing at the cell edge (Figure 5D). Instead, mCIT-2xPH^PLC^ accumulated in the cytosol, while still being excluded from the round central nuclei (Figure 5E). Note that after dex induction, *MAP-mCH-dOCRL* was strongly expressed in the peripheral zone, and was only weekly expressed in the central zone (Figure 5E). However, although its expression was weak in the central zone it appeared to be sufficient to impact mCIT-2xPH^PLC^ localization (Figure 5E). Taken together, our results indicate that iDePP was efficient for PI(4,5)P_2_ depletion in both NPA-treated and dissected shoot apical meristems (Figure 5C, D; Supplemental file 4). However, because iDePP is not uniformly induced in this tissue, the expression of *MAP-mCH-dOCRL* or its impact on mCIT-2xPH^PLC^ localization should always be double checked before drawing any conclusions.

Next, we analyzed the impact of iDePP in the epidermis of cotyledons. Most notably, we focused on the pavement cells, an epidermal cell with a jigsaw-like shape. Pavement cells are differentiated cells with a big central vacuole that pushes the cytoplasm against the plasma membrane and the cell wall. Thus, the fluorescent protein fusions that are localized in the cytosol in this is cell type are enriched at the cell periphery. However, unlike plasma membrane targeted fluorescent fusion proteins, which have a sharp localization at the cell periphery, the localization of cytosolic proteins is more diffuse and they are also present in cytoplasmic strands. In the absence of dex, MAP-mCH-dOCRL was not expressed and mCIT-2xPH^PLC^ was sharply localized at the cell periphery without the presence of cytoplasmic strands (Figure 5E), likely being at the plasma membrane. Dex induction of MAP-mCH-dOCRL was efficient in this cell type and impacted the localization of mCIT-2xPH^PLC^, which became wider at the cell edge and also localized to cytoplasmic strands (Figure 5E). This localization indicated that mCIT-2xPH^PLC^ was solubilized upon *MAP-mCH-dOCRL* induction in pavement cells and suggested that PI(4,5)P_2_ is hydrolyzed in this condition. Similar to the situation in roots, the *MAP-mCH-dOCRL*^*dead*^ and *MAP-3xmCH* controls were efficiently expressed in this cell type upon dex induction, but they had no effect on mCIT-2xPH^PLC^ localization, which remained at the plasma membrane and was not found in the cytoplasm (Figure 5F and G).

Taken together, these results suggest that our iDePP system efficiently depletes PI(4,5)P_2_ in various cell types and organs, both in roots and aerial parts.

### Biological relevance: iDePP-mediated PI(4,5)P_2_ depletion affects microtubule organization in elongating root epidermal cells

Once the inducible PI(4,5)P_2_ depletion established, we next set-out to test the cellular effects of PI(4,5)P_2_ depletion on actin organization in plants, since this lipid is well known as a master regulator of actin dynamics in other eukaryotes (Logan and Mandato, 2006; Moss, 2012; Zhao et al., 2010). For example, depletion of PI(4,5)P_2_ at the plasma membrane by expressing phosphatases disrupts the actin cytoskeleton leading to a cell-wide reduction in polymerized F-actin in animal cells (Raucher et al., 2000; Shibasaki et al., 1997). Since the activity of actin-remodeling protein are known to be regulated by PI(4,5)P_2_ in plant cell (Gungabissoon et al., 1998; Guo et al., 2019; Huang et al., 2003; Staiger et al., 1997), we investigated qualitatively whether PI(4,5)P_2_ depletion could affect the organization of F-actin cytoskeleton in plant cells in a manner similar to that of animal cells. Using live cell imaging, we observed actin filament in elongated root cell using z-projection of the F-actin marker LifeAct-YFPv before and upon PI(4,5)P_2_ depletion using iDePP. Surprisingly, expression of MAP-mCH-dOCRL, did not led to a reduction in polymerized F-actin in epidermal root cells or to any major rearrangement of F-actin cables, at least qualitatively, compared with of the MAP-3xmCH control (Figure 6A).

**Figure 6.**
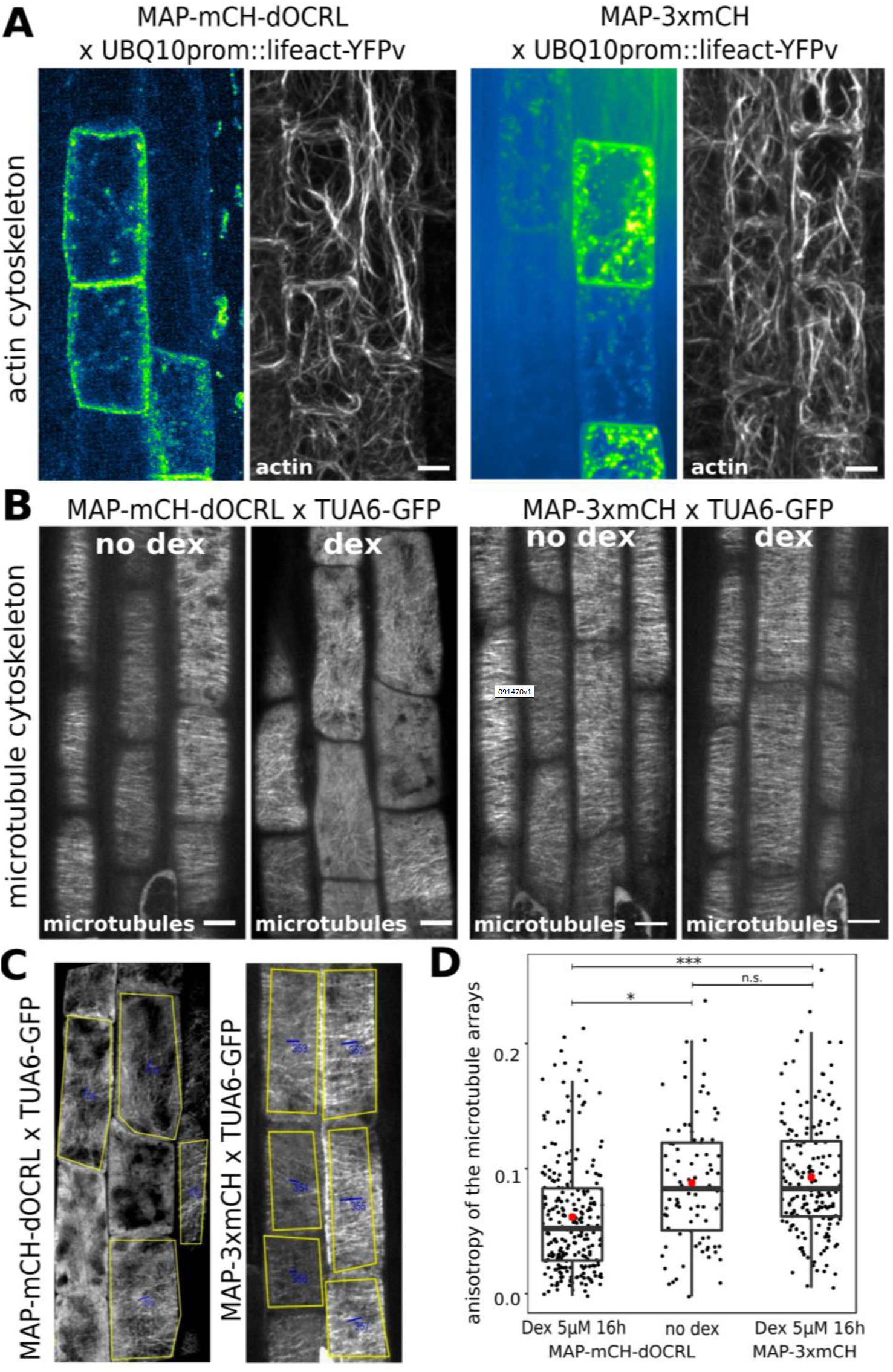
Effect of PI(4,5)P_2_ depletion on cytoskeleton dynamics in epidermal root cells. (A) F-actin cytoskeleton labelled by LifeAct-YFPv in cells expressing MAP-mCH-dOCRL or MAP-3xmCH. (B) Cortical microtubules labelled by TUA6-GFP, in non-treated cells (no dex) or treated (dex for 16 hours) to induce MAP-mCH-dOCRL or MAP-3xmCH expression. All pictures are z-projections. no dex: no treatment. dex: dexamethasone treatment. Scale bars: 10 µm. (C) Example of the images used for the quantification of the anisotropy using FibrilTool macro for imageJ. (D) Average anisotropy of the microtubule arrays in MAP-mCH-dOCRL or MAP-3xmCH lines. For statistical analysis, we used a linear mixed effect model and then computed post hoc tests (*lsmeans* package). Details for the statistical analysis can be found in Supplemental Table 5. ***: p-value < 0.0005; **: p-value < 0.05; n. s.: p-value ≥ 0.05.

We next investigated a possible role of PI(4,5)P_2_ on the organization of cortical microtubules in *Arabidopsis* elongated root cells, using TUBULINα6-GFP (TUA6-GFP) protein fusion. In the root elongation zone, the cortical microtubules of interphasic epidermal cells are transversely aligned (Vaughn et al., 2011), thereby forming a network that gates unidirectional anisotropic cell expansion. In absence of dex induction in *MAP-mCH-dOCRL* line or upon expression of *MAP-3xmCH*, cortical microtubules of elongating epidermal cells were observed to form a network of transversely aligned microtubules, orthogonal to the elongation axis of the cells as expected (Figure 6B, Polko and Kieber, 2019). This result was quantitatively reflected by a relatively high anisotropy of the microtubule array per cell (mean ± SEM is 0.095 ± 0.005 for the MAP-mCH-dOCRL no dex, n = 90 cells and mean ± SEM is 0.099 ± 0.003 for the MAP-3xmCH with dex, n = 175 cells; Type II Wald χ^2^ test: χ^2^ = 12.155, p = 0.9387; Figure 6D, Supplemental file 5). Depletion of PI(4,5)P_2_ after induction of MAP-mCH-dOCRL led to the increased randomization of cortical microtubule orientations, which correlates with an increase in isotropy of microtubule arrays (mean ± SEM is 0.067 ± 0.003 for the MAP-mCH-dOCRL with dex, n = 238 cells, Type II Wald χ^2^ test: χ^2^ = 12.155, p = 0.0047, Figure 6C, D, Supplemental file 5). This result suggests that cortical microtubule reorganization requires PI(4,5)P_2_ at the plasma membrane at least during the elongation process of epidermal root cells, and validate our approach to study the role of PI(4,5)P_2_ in a time-scale compatible with plant cell morphodynamics.

## CONCLUSION AND DISCUSSION

Here, we describe iDePP, a new genetically encoded system that allows to rapidly deplete PI(4,5)P_2_ from the plasma membrane *in planta*. The stable genetically encoded system we describe here is unprecedented and allow to specifically induce PI(4,5)P_2_ depletion *in planta*. It is also one of the first in any multicellular organism (Guglielmi et al., 2015). We provide *in vitro* evidence that *Drosophila melanogaster* dOCRL phosphatase domain can specifically dephosphosphorylates PI(4,5)P_2_. The use of adequate negative controls demonstrates that the observed effect on PI(4,5)P_2_ sensor only depends on dOCRL enzymatic activity and is not caused by side effects of the dex inducible system or the recruitment of an endogenous PI(4,5)P_2_ phosphatase. Also, the absence of strong effect observed on various other membrane lipids, including PI4P, PS, and PI3P, indicated that the system specifically targets PI(4,5)P_2_ without massively affecting the plasma membrane and internal membranes. Yet, our results suggest link between PI(4,5)P_2_ and PS accumulation in plants, as it was recently suggested in animal systems (Ghai et al., 2017; Sohn et al., 2018).

MAP-mCH-dOCRL fluorescence was detected as soon as three hours—sometimes even earlier— after gene induction was triggered. This roughly corresponds to the time required for readily detection of GFP-HsPIP5KIα after oestrogen treatment in (Gujas et al., 2017) system that increases PI(4,5)P_2_ levels. Interestingly, we detected solubilization of the mCIT-2xPH^PLC^ PI(4,5)P_2_ sensor before we could detect MAP-mCH-dOCRL. This correlation strongly suggests that MAP-mCH-dOCRL is highly efficient and strongly depletes PI(4,5)P_2_ even when its concentrations are below fluorescence detection levels or prior to the full maturation of the fluorescent reporter mCH. The system allows to rapidly — couple of hours— deplete PI(4,5)P_2_, thereby circumventing the long term effects that a chronic PI(4,5)P_2_ depletion would have, for example through a constitutive overexpression of a plasma membrane-targeted 5-phosphatase or genetic elimination of key enzyme responsible for PI(4,5)P_2_ homeostasis (e.g. PIP5Ks).

Importantly, solubilization of mCIT-2xPH^PLC^ PI(4,5)P_2_ sensor was effective in both root and shoot tissues. We can even speculate that in other cell types such as pollen tubes, this system will be relevant to study the role of the patterning of PI(4,5)P_2_ in polarized growth. Here, we focused on epidermal tissues, because they are relatively easier to image at the same time the induction of MAP-mCH-dOCRL and the anionic lipid biosensors. However, MAP-mCH-dOCRL is induced in the broad expression domain of *UBQ10* promoter, and thus iDePP may be used in additional tissues and organs. Alternatively, MAP-mCH-dOCRL can easily be expressed in a tissue specific manner (Marquès-Bueno et al., 2016). The spatial control of PI(4,5)P_2_ depletion, even if not reversible, would likely be of great help to elucidate PI(4,5)P_2_ functions in cell differentiation and plant development.

We observed that expression of *MAP-mCH-dOCRL* led to defects in cortical microtubules orientation, which often failed to adopt a proper anisotropic organization orthogonal to cell growth direction in elongating root epidermal cells. This observation suggests that the maintenance of anisotropic-microtubules arrays in elongated cells requires PI(4,5)P_2_. This may affect proper cellulose microfibrils deposition by microtubule-guided cellulose synthase complexes. Phospholipids have been involved in the control of cytoskeleton cortical anchoring and dynamics both in plants and animals (Raucher et al., 2000; Zhang et al., 2012). The increased isotropy of microtubule arrays and the slower response to mechanical perturbations in mutants impaired in microtubule self-organization correlate with an altered phosphoinositide patterning within the shoot apical meristem (Colin and Jaillais, 2020; Stanislas et al., 2018). Indeed, the katanin mutant allele *botero1-7* exhibits more disorganized microtubules, reduced growth anisotropy and slower microtubule response to mechanical perturbations (Bichet et al., 2001; Lindeboom et al., 2013; Uyttewaal et al., 2012) and the PI(4,5)P_2_ pattern is noisier in *bot1-7* than in the wild type (Stanislas et al., 2018). This results, together with our finding that the perturbation of the PI(4,5)P_2_ patterning affects microtubules anisotropy suggest that PI(4,5)P_2_ at the plasma membrane promotes microtubule array order. It was proposed that KTN1 function through RHO-LIKE GTPASE FROM PLANTS 6 (ROP6) and its effector ROP-INTERACTIVE CRIB MOTIF-CONTAINING PROTEIN 1 (RIC1), which binds to and activates KTN1 (Fu et al., 2009; Lin et al., 2013; Xu et al., 2010). ROP6 requires phosphatidylserine for signaling at the plasma membrane but it also interact with PI(4,5)P_2_ in vitro (Platre et al., 2019). Future directions should thus include the characterization of the molecular player responsible for the PI(4,5)P_2_ accumulation and signaling at the plasma membrane and their role in the remodeling of the microtubule cytoskeleton leading to directional growth.

The iDePP system should prove useful for the community of plant biologists and may be amended, by using tissue specific promoters for instance. Also, instead of dOCRL 5-phosphatase domain, others domains with other enzymatic activity could be similarly targeted at the plasma membrane. Such domains could allow depleting other species of plasma membrane lipids, but also constitute systems to assess the enzymatic activities of putative PI4P and/or PI(4,5)P_2_ phosphatases and phospholipases *in vivo*. iDePP can also be used to assess a protein requirement to PI(4,5)P_2_ for plasma membrane targeting. Furthermore, the timing of iDePP activity is fully compatible with in vivo live imaging. Together, we believe that the iDePP system will have the potential to be useful for numerous research groups working in the fields of plant cell biology, plant development but also immunity and reproduction.

## MATERIAL AND METHODS

### Sequence alignment

HsOCRL_234-539_ and dOCRL amino acid sequences were aligned using T-Coffee software (http://tcoffee.crg.cat/apps/tcoffee/do:regular) and the fasta_aln files obtained were then treated with Boxshade (http://www.ch.embnet.org/software/BOX_form.html) and inkscape programs.

### Cloning

A synthetic gene (https://eu.idtdna.com/pages) corresponding to dOCRL_168-159_ codon optimized for expression in *Arabidopsis thaliana* (see sequence below) was amplified and flanked with NcoI and XhoI restriction sites by PCR and cloned into pMH-HS-SUMO vectors (Christie et al., 2012) by restriction and ligation (see corresponding primers in Supplemental Figure 6).

dOCRL_168-159_:

gtaaagcaagagttaaagaagcgagagagtgagtatatcgtctataaggacattatcatttactgcgctacgtggaacgtgaacaataagacatgcagtgatagtaacaatccgctaagagcgtggctcgcatgctccgaaaagccgccagacatatatgcgataggtcttcaggagctagacaccccgaccaaggctatgctcaatagtacccaagtccaagcaattgaaaaacagtggatagataagatgatggacagtgtgcatcctgacgtagagtacgagattttgatgtctcaccgattagttgcgacgatgcttactgtcatcgttagaaaacagcttaggcagcacattatacgttgtcgacccaaatccgtggcccgtggaatattcaacacgctcggaaataagggaggggtggcaatatccctgcagttgaatgagggaaacatatgttttgttaactcccatctggctgcacatatggggtacgtcgaagagaggaatcaagattacaacgctattgtcgaaggcatcaggtttgacgacggtagaactatctccgaccatgaccatatattttgggtaggagacttgaattatcgaatacaggaacctccgggacagcagcgtcctgggccgttgagtgacgcacagacatacgagctcttattacaatacgatcaactccgtcaagagatgcgaaggggaaaatgcttcgaaggttacacggagggggagattaagttcagaccgacatacaaatatgatcctggaacagacaattatgactcttcagagaaacagcgagcacctgcatactgcgatagagtgctatggaaaggtacacgtatcgaacagctagcatacaacagcataatggaaattaggcaaagcgatcataagccagtttatgcggttttccaggtgaaggtaaagacacgagatgaggtgaagtataagcgtgttcaagaggaggtgctaaaggcggttgacaaaagggagaacgataatcagccacagtaa

*UBQ10pro:GVG* (*UBQ10pro:GVG-tE9::6xUAS-35Smini::* rigorously) construct was cloned using Gibson technique and cloned into a *pDONR-P4P1R* gateway vector (thermofisher cat# 12537023). *MAP-mCH* was amplified from a *mCH/pDONR207* matrix using 5’ phosphorylated primers and ligated into *MAP-mCH/pDONR207*. The synthetic dOCRL_168-509_ sequence (see above) with an N-terminal GAGARS linker (ggggcaggagccagatcc) was flanked with attB2R and attB3 sequences and recombined by BP gateway reaction into *pDONR-P2RP3*. Site directed mutagenesis of dOCRL was performed by PCR on *dOCRL*_*168-509*_/*pDONR-P2RP3* using 5’ phosphorylated primers, and subsequent ligation of the linear plasmid circularized *dOCRL*_*168-509*_^*D468G*^/*pDONR-P2RP3*. 2xmCHERRY/*pDONR-P2RP3* was already available.

*UBQ10pro:GVG:MAP-mCH-dOCRL*_*168-509*_*/pH7m34GW, UBQ10pro:GVG:MAP-mCH-dOCRL*_*168-509*_^*D468G*^*/pH7m34GW*, and *UBQ10pro:GVG:MAP-3xmCH//pH7m34GW* were obtained by triple LR gateway reaction between *UBQ10pro:pDONR-P4P1R, MAP-mCH-pDONR207*, the corresponding *pDONR-P2RP3* vectors, and *pH7m34GW* destination vector (Karimi et al., 2007). See resource table for details.

### Protein purification

BL21(DE3) pLys competent *E. coli* (Novagen and Supplementary file 5) were transformed with *pMH-HS-Sumo-dOCRL*_*168-509*_ vector (Kan^R^) and cultured on Luria Broth (LB) liquid or 1 % agar medium supplemented with 50 µM Kanamycin. Two 800 ml 2xTY (10 g/L yeast extract; 16 g/L tryptone; 5 g/L NaCl) liquid cultures supplemented with kanamycin were inoculated with 20 ml from an overnight pre-culture of 150 ml, and left at 37°C with 200 rpm shaking. When OD_600nm_ reached 0.6, (i) protein expression was induced by adding IPTG 0.625 mM final concentration, (ii) 45 ml of glycerol were added, and (iii) temperature was lowered by transferring the liquid culture on ice for 15 min. The induced culture was then grown for 18 h at 18°C with 200 rpm shaking. Bacteria were then centrifuged 30 min at 6000 g and 4 °C, and bacterial pellets scooped, resuspended in lysis buffer (200 mM NaCl; 20 mM Tris-HCl pH 7.5; 1 mM PMSF) with the aid of vortex, flash frozen and stored at −80°C. The day of protein purification, Bacterial pellets corresponding to 33 % (270 ml of the culture) were thawed at room temperature. For then on, all steps were carried on ice. Bacteria were then sonicated in four pulses of 30 sec and 30 W. Sonicated lysates were centrifuge 30 min at 13000 g and 4°C in new falcon tubes.

Supernatants, corresponding to the soluble fraction of the lysate, were then diluted to 50 ml final volume and subjected to affinity chromatography. 2 ml of 50 % HIS-Select® Cobalt affinity resin (Sigma) was washed with milli-Q water and then with equilibration buffer (20 mM imidazole; 200 mM NaCl; 20 mM Tris-HCl pH 7.5; 1 mM). Bacterial lysate was run thought the resin twice. Then, resin was washed with wash buffer (40 mM imidazole; 200 mM NaCl; 20 mM Tris-HCl pH 7.5; 1 mM).

Elution was performed with three fractions of 2 ml of elution buffer (250 mM imidazole; 200 mM NaCl; 20 mM Tris-HCl pH 7.5; 1 mM). Elution fractions were then either directly concentrated (C1 fraction) or kept at 4 °C for a maximum of 16 h before concentration (C2 fraction). Concentration was achieved using 0.5 ml 30 kD molecular-weight cut-off concentration columns (Millipore UFC 501024) centrifuged at 4000 g and 4 °C until 2 ml of elution reached a final volume of 400 µL. Aliquots of 20 µL were sampled from induced bacterial culture (IN), flow thought (FT), wash (W), elution fraction (E1, E2 and E3) and concentrated fractions (C1 and C2). They were mix with SDS loading buffer (0.0625 M Tris HCl pH 6.8; 2.5 % SDS; 2 % DTT (0.13 M); 10 % glycerol; bromophenol blue), vortexed and submitted for 5 min to 95°C treatment, and then stored at −20°C. Protein concentration in concentrated fraction was calculated from absorbance at 280 and 260 nm measured with a nanodrop and calculated extinction coefficient (https://web.expasy.org/protparam/). His-Sumo-dOCRL_168-509_-Hishas an expected molecular weight of 54.82 kD.

### Malachite green phosphatase assay

Phosphate released was monitored using a Malachite green phosphatase kit (Echelon) and water soluble short chain (di:C8) phosphoinositides (Echelon) stored in glass vials at −20°C as stock solutions of 500 µM in CHCl3/MetOH (9:1; PI, PI3P, PI4P, PI5P), or in CHCL3/MetOH/H2O (5:5:1; PI(3,5)P_2_, PI(4,5)P_2_). di:C16 Phosphatidylserine (PS; Echelon) was stored in glass vials at −20°C as a stock solution of 10 mM in CHCl3/MetOH (9:1). Before malachite green assay, inorganic solvent was evaporated by centrifugation in a speed vac at 45 °C for 15 min. Dried lipids (PS and one phosphoinositide) were then resuspended in reaction buffer (200 mM NaCl 50 mM Tris-HCl pH 7.5; 10 mM KCl; 5 mM MgCl_2_) with a 30 sec of sonication at full power. Reaction were carried out in a 20 µL volume consisting in reaction buffer containing 500 µM of PS, 100 µM of a given short chain phosphoinositide species (except for the “mock” condition), and 5 µL of concentrated protein fractions (C1 or C2; except of “no enzyme” condition) corresponding to the final concentration displayed on Figure 1C. In the “iP” condition corresponds to addition of inorganic phosphate. Malachite green assay was carried out at 29 °C for 75 min, with gentle shaking every 15 min, and reaction was eventually quenched by the addition of 90 µL of malachite green reagent. Pictures were taken 25 min after the addition of Malachite green reagent, and green coloration qualitatively indicated a significant phosphate release. This Experiment was carried out independently three times, with either one or two replicates each time, and gave consistent results.

### Growth condition and plant materials

*Arabidopsis thaliana* Columbia-0 (Col-0) accession was used as wild type (WT) reference genomic background throughout this study. *Arabidopsis thaliana* seedlings *in vitro* on half Murashige and Skoog (½ MS) Basal Medium supplemented with 0.8% plant agar (pH 5.7) in continuous light conditions at 21 °C. Plants were grown in soil under long-day conditions at 21 °C and 70% humidity 16 h daylight.

### Plant transformation and Selection

Plant were transformed and selected as previously described (Platre et al., 2018). Each construct was transformed into *Agrobacterium tumefaciens C58 GV3101* strain and selected on YEB media (5g/L beef extract; 1g/L yeast extract; 5g/L peptone; 5g/L sucrose; 15g/L bactoagar; pH 7.2) supplemented with antibiotics (Spectinomycin, Gentamycin, Rifampicin). After two days of growth at 28 °C, bacteria were scooped and re-suspended in roughly 200 mL of transformation buffer (10mM MgCl_2_; 5% sucrose; 0.25% silweet) and Col-0 *Arabidopsis* were transformed by dipping. Primary transformants (T1) were selected *in vitro* on the appropriate antibiotic/herbicide (glufosinate for mCIT, hygromycin for mCH-tagged proteins). Approximately 20 independent T1s were selected for each line. In the T2 generation at least two independent transgenic lines were selected using the following criteria when possible: (i) good expression level in the root for detection by confocal microscopy upon dexametazone induction, (ii) relatively uniform expression pattern, (iii) line with no obvious abnormal developmental phenotypes. Lines were rescreened in T3 using similar criteria as in T2 with the exception that we selected homozygous lines (100% resistant). At this step, we selected one transgenic line for each marker that was used for further analyses and crosses. NPA treatments were carried out as previously described (Grandjean et al., 2004). Plants were further kept on NPA-containing medium for all the duration of the experiments.

### Induction of gene expression with dexamethasone

In order to study the effect of iDePP system in root, seedlings of each genotype were grown vertically for five to seven-day *in vitro* on half Murashige and Skoog (½ MS) Basal Medium supplemented with 0.8% plant agar (pH 5.7) in continuous light conditions at 21 °C. Part of the seedlings were next carefully transferred to ½ MS medium plates, 0.8 % agar (pH 5.7), containing 5 µM dexamethasone. Both non-treated seedlings and dex-treated seedlings were, all together, grown back in continuous daylight (see before). 16 h after transfer on dexamethasone plates, (more or less 90 min) seedlings were mounted in ½ MS medium and imaged by confocal microscopy.

For cotyledon analysis, seeds were sterilized and sown on Arabidopsis medium (MS medium without sugars and vitamins) for 7days. Seedling were then carefully transferred on MS medium (without sugars and vitamins) supplemented with 5µM of Dexamethasone. Cotyledons were placed at the surface of the medium, as flat as possible, while the rest of the plant was immerged into the medium.

Shoot apex were dissected as described by (Stanislas et al., 2017) and then placed in a shoot apex medium as described by (Grandjean et al., 2004) supplemented with 5µM of Dexamethasone. To generate naked meristems in vitro, seeds were first directly sown on Arabidopsis medium, supplemented with 10 μM NPA. Seedlings with naked meristems were then selected and transferred to a shoot apex medium without NPA and supplemented with 5µM of Dexamethasone. Images were taken at 18 h after induction.

### Time-lapse imaging

Time lapse imaging of cell division and root hair growth were performed mostly as described previously (Doumane et al., 2017), with few amendments. In brief, five days old Arabidopsis seedlings were transferred in a chambered cover glass (Lab-Tek II, www.thermoscientific.com), which contained 3 ml of ½ MS medium (pH 5.7) containing 0.8 % plant agar (Duchefa, http://www.duchefa-biochemie.nl/) and 5 µM dexamethasone and (ii) 800 µL of ½ MS medium (pH 5.7) containing 5 µM dexamethasone. Epidermal cells in the meristematic region of the root tip were subjected to time-lapse imaging with spinning disk confocal microscope. Up to three roots were observed simultaneously and images were collected at different Z-positions every 5 min for 8 h. Tracking of growing roots was automatically performed (Doumane et al., 2017).

### Microscopy setup

Images of cotyledons, shoot apical meristem and NPA meristems (Figure 4B-D) were acquired with a Leica SP8 upright scanning confocal microscope equipped with a water immersion objective (HCX IRAPO L 25x/ 0.95 W). Fluorophores were excited using Led laser (Leica Microsystems, Wetzlar, Germany) emitting at wavelengths of 514 nm for mCIT fluorochrome and 552 nm for mCH fluorochrome. Emission fluorescence was recovered between 519nm - 548nm for mCIT and between 604 nm - 662 nm for mCH. The following scanning settings were used: pinhole size 1 AE, scanning speed of 8000 Hz (resonant scanner), frame averaging between 8 and 10, Z intervals of 0.45 μm.

All additional imaging was performed with the following spinning disk confocal microscope set up: inverted Zeiss microscope (AxioObserver Z1, Carl Zeiss Group, http://www.zeiss.com/) equipped with a spinning disk module (CSU-W1-T3, Yokogawa, www.yokogawa.com) and a ProEM+ 1024B camera (Princeton Instrument, http://www.princetoninstruments.com/) or Camera Prime 95B (www.photometrics.com) using a 63x Plan-Apochromat objective (numerical aperture 1.4, oil immersion). GFP and mCIT were excited with a 488 nm laser (150 mW) and fluorescence emission was filtered by a 525/50 nm BrightLine! single-band bandpass filter (Semrock, http://www.semrock.com/), mCH was excited with a 561 nm laser (80 mW) and fluorescence emission was filtered by a 609/54 nm BrightLine! single-band bandpass filter (Semrock, http://www.semrock.com/). For quantitative imaging, pictures of epidermal root meristem cells were taken with detector settings optimized for low background and no pixel saturation. Care was taken to use similar confocal settings when comparing fluorescence intensity or for quantification. Signal intensity was color-coded (green fire blue scale, https://fiji.sc/).

### 3D projections, dissociation indexes and anisotropy

The fluorescence intensity in root hair was obtained using Fiji. A line of 66 pixels was drawn at the proximity of the root hair tip and the intensity of grey was plot using the Plot Profil tool. The values were transferred in excel file to obtain the graph. Shoot apical meristem and NPA-induced meristem projected images were obtained by using the MorphoGraphX software (https://www.mpipz.mpg.de/MorphoGraphX). The signal projections were generated by extracting the fluorescent signal at the surface of the meristem (between 2 µm and 5 µm from the meristem surface) and by projecting it on the cellular mesh. Projection of root epidermal cells were obtained using Fiji “Max intensity projection” tool on Z-stacks of 21 slices distant of 0.5 µm from each other.

Dissociation indexes of membrane lipid fluorescent biosensors were measured and calculated as previously described (Platre et al., 2018). Briefly, we calculated “indexNoDex” in the mock condition, defined as the ratio between the fluorescence intensity (Mean Grey Value function of Fiji software) measured in two elliptical regions of interest (ROIs) from the plasma membrane region (one at the apical/basal plasma membrane region and one in the lateral plasma membrane region) and two elliptical ROIs in the cytosol. Next, we measured a similar ratio after dexamethasone treatments (‘‘indexDex’’). The dissociation index is the ratio of (indexNoDex)/(indexDex). This dissociation index reveals the degree of relocalization of the fluorescent reporters from the plasma membrane to the cytosol, between the non-treated and perturbed conditions (pharmacological treatment or mutant). Dissociation indexes of mCIT-2xPH^PLC^ during time-lapse induction of MAP-mCH-dOCRL (Figure 4) were measured in 30 cells of the same root at each time point.

For quantification of the anisotropy of microtubule arrays in the different transgenic lines, Maximal z projection of z-stack of epidermal root cell in the elongation zone were obtained, using Fiji. In fibriltool, the value for ‘Multiply line length by’ was set up at 1 and ROI using the built-in Polygon tool were generated for individual cells (>100 cells per conditions). The data from the log file was used to extract the average anisotropy of microtubule arrays (a score between 0 and 1) and to run statistical analyses.

### Statistical analysis

For dissociation index and anisotropy, we performed all our statistical analyses in R (v. 3.6.1, (R Core Team, 2019), using R studio interface and the packages ggplot2 (Wickham, 2016), lme4 (Bates et al., 2015), car (Fox and Weisberg, 2011), multcomp (Hothorn et al., 2008) and lsmeans (Lenth, 2016). For each model assuming normality, we plotted residuals values against normal quantiles to confirm their normal distribution. Graphs were obtained with R and R-studio software, and customized with Inkscape (https://inkscape.org).

## ACKNOLEDGMENTS

We are grateful to the SiCE group (RDP, Lyon, France) in particular Thierry Gaude and Vincent Bayle (RDP, Lyon), and to Yohann Boutté (LBM, Bordeaux, France), Daniël Van Damme (VIB, Ghent, Belgium), Thomas Stanislas (ZMBP, Tübingen, Germany), Fabrice Besnard and Nicolas Doll (RDP, Lyon, France), for comments and discussions. We as well thank former interns Lucas Courgeon (ENS de Lyon) and Amélie Bauer (UCB Lyon 1 University), who performed preliminary experiments during early steps of this project and helped with plant labor, and Karin Grünwald. We thank Justin Berger, Patrice Bolland, and Alexis Lacroix from our plant facility, and Claire Lionnet from the imaging platform. We also acknowledge precious help from Romain Boisseau (OBEE department, University of Montana, USA) regarding the statistical analysis. This work was supported by ERC no. 3363360-APPL under FP/2007–2013 (YJ), ANR caLIPSO (ANR-18-CE13-0025-02; YJ), ANRJC/JC JUNIOR INVESTIGATOR GRANT (ANR-16-CE13-0021; MCC, AF) and a SEED FUND ENS LYON-2016 (MCC). MD, LC and AL are funded by Ph.D. fellowships from the French Ministry of Research and Higher Education.

## ADDITIONAL FILES

**Supplementary file 1.**
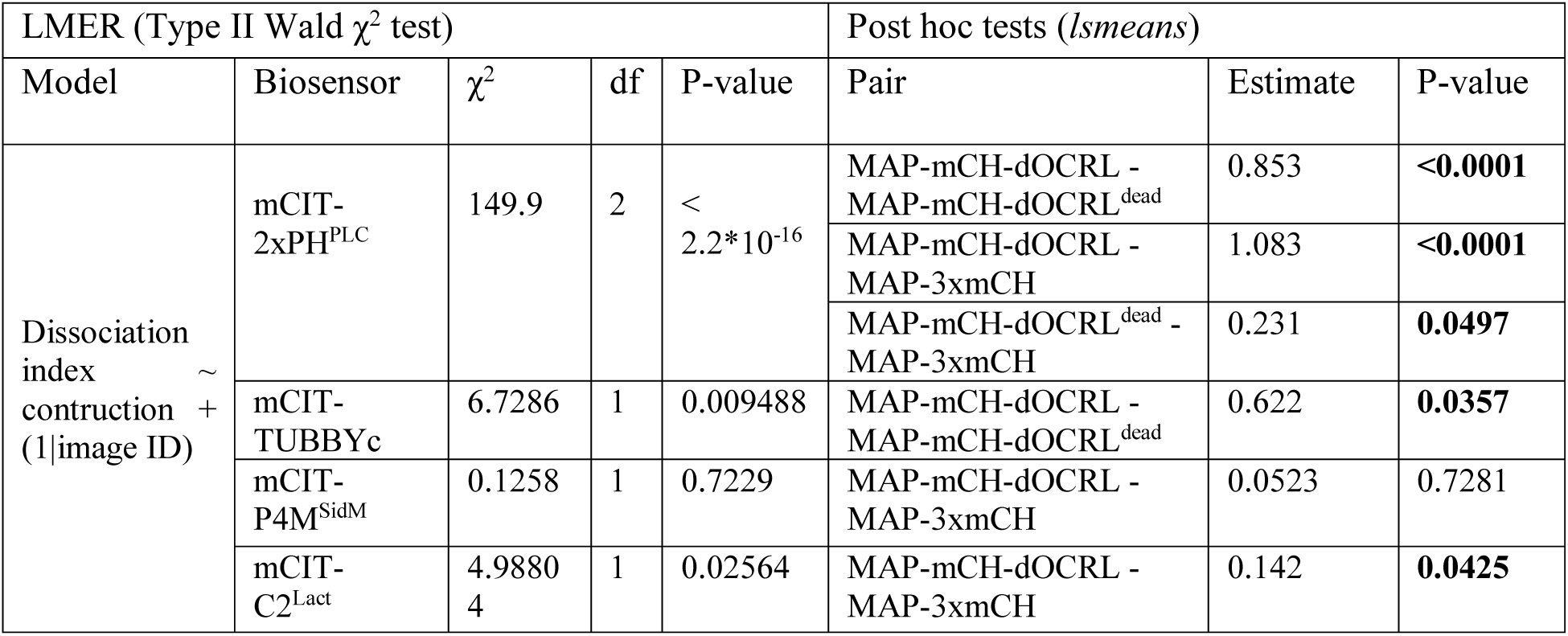
Statistical analysis of lipid biosensor dissociation indexes. For various plasma membrane localized lipid biosensors, we compared dissociation indexes across inductions of constructions encoding MAP-mCH-dOCRL or negative controls (MAP-mCH-dOCRL^dead^ and MAP-3xmCH). We used a linear mixed effect model accounting for image ID (*id* est root) as random factor (as one root per image were imaged, but ten cells per root used). We then computed post hoc tests (*lsmeans* package)

**Supplementary file 2.**
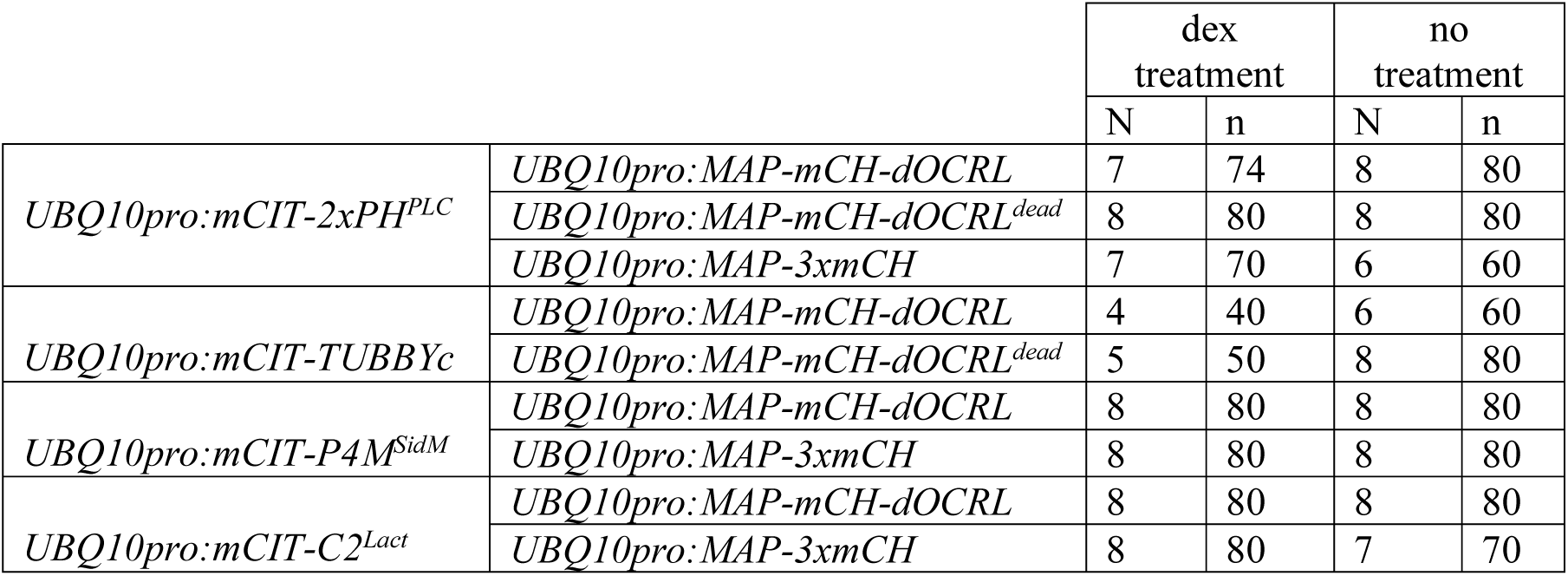
Size of workforces corresponding to dissociation indexes calculations. N: number of roots; n: number of cells (10 cells per roots except for 14 cells taken in one root (first line)).

**Supplemental file 3.**
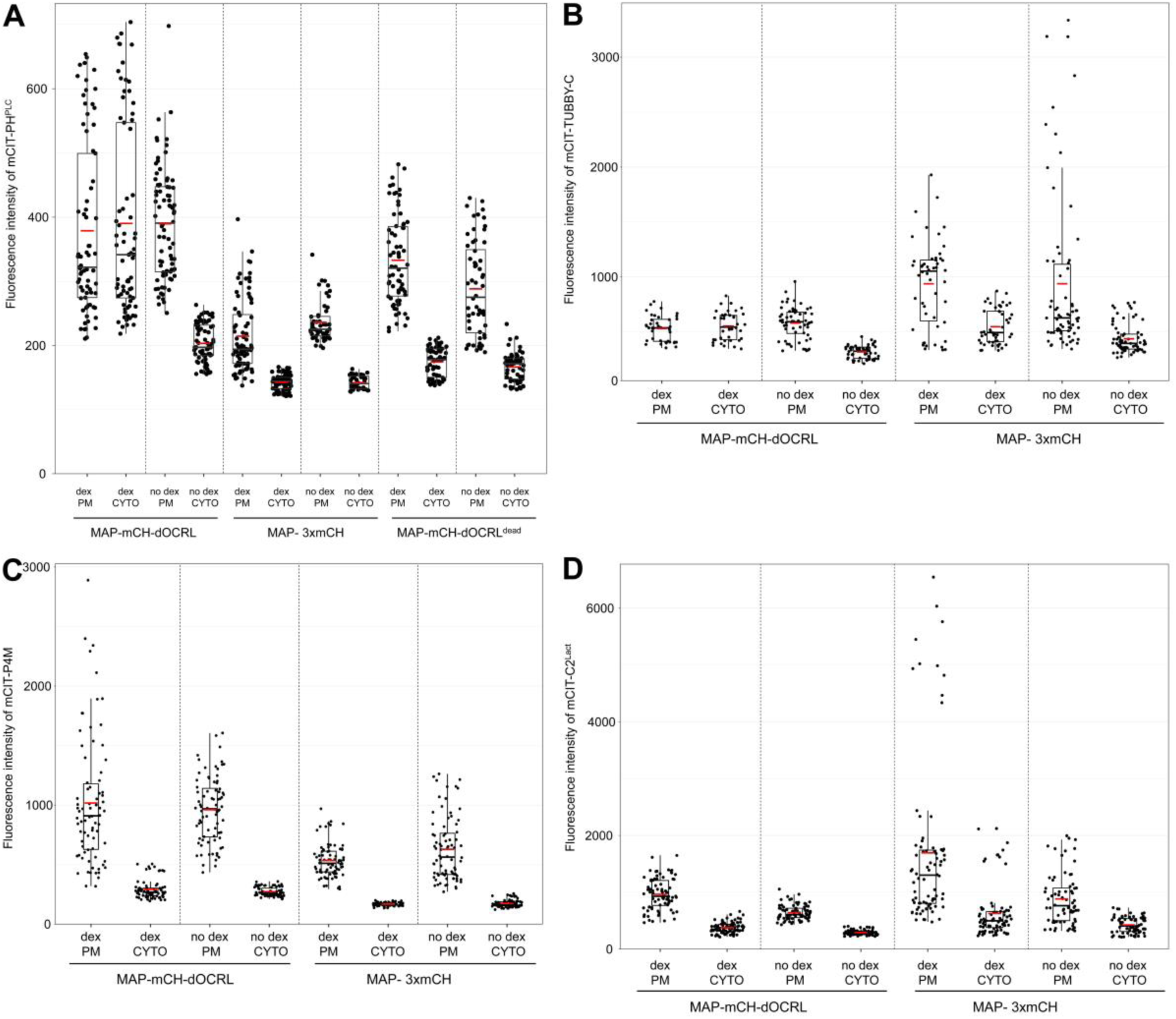
Graphic representation of the fluorescent intensity observed for the different biosensors, used for the quantification of the dissociation index presented in Figure 3G.

**Supplemental file 4.**
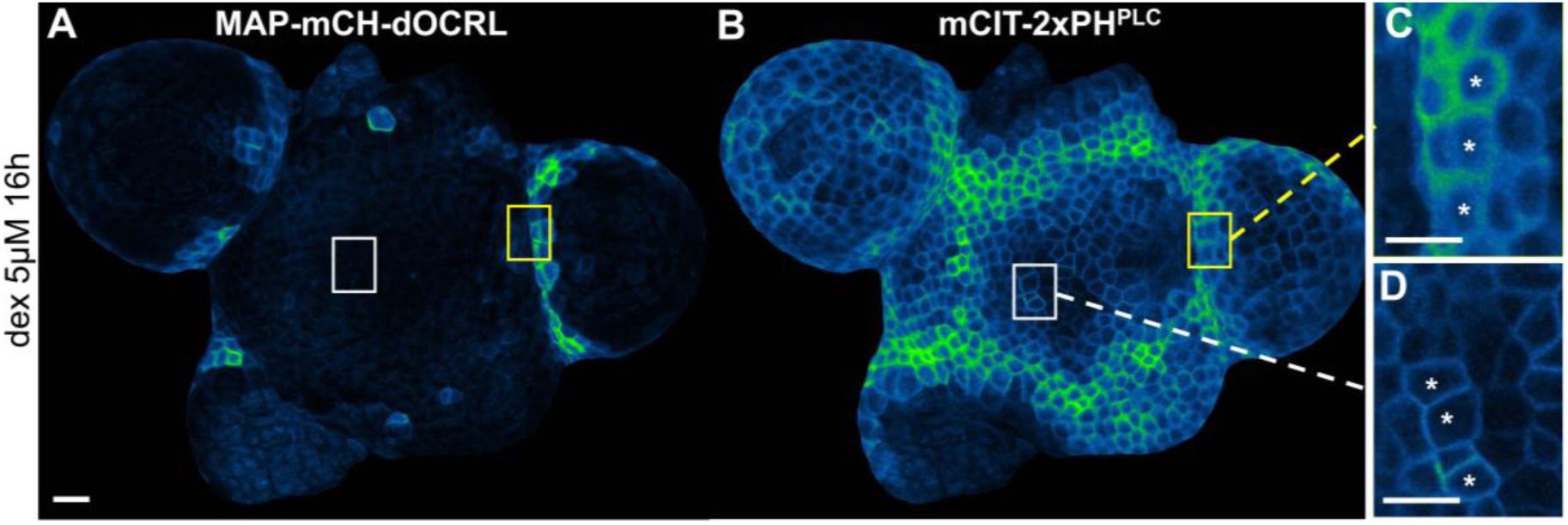
Induction of the iDePP system in the shoot apical meristem of Arabidopsis. (A) Subcellular localization of MAP-mCH-dOCRL after dex induction. (B) Subcellular localization of mCIT-2xPH^PLC^ in the same shoot apical meristem after dex induction. (C-D) Close-ups from (B). The yellow ROI in (A), (B) and (C) represents the fluorescence observed in a zone where MAP-mCH-dOCRL was not induced. The white ROI in (A), (B) and (D) represents the fluorescence observed in a zone where MAP-mCH-dOCRL was induced. Scale bar, 20 μm in (A-B), 10 μm in (C-D). Asterisks represent the position of the nucleus in the cell.

**Supplementary file 5.**
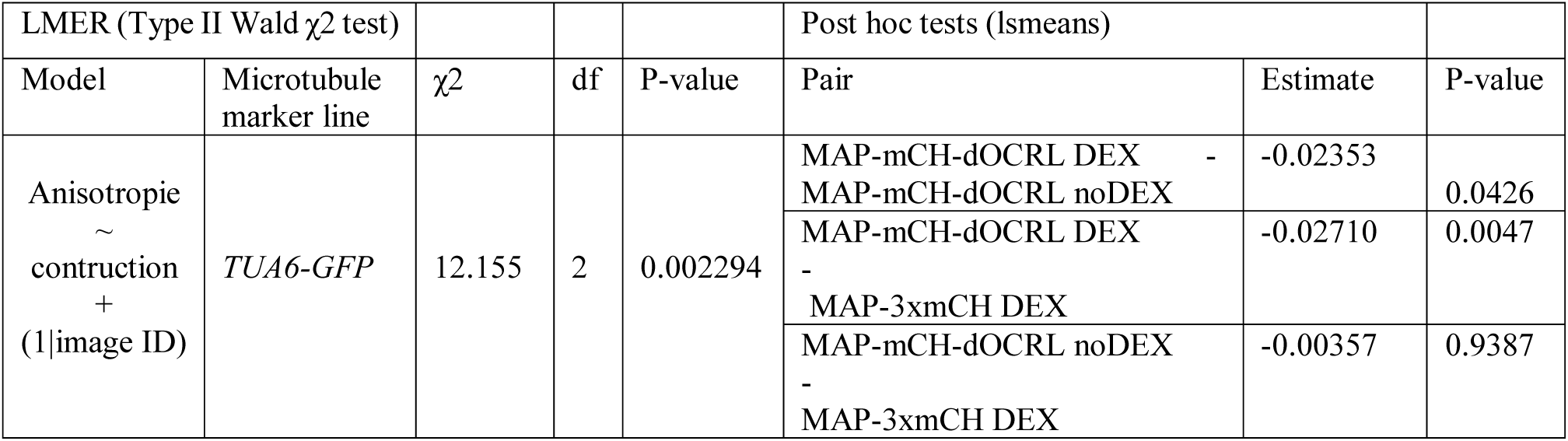
Statistical analysis of microtubules anisotropy per cells. We used a linear mixed effect model accounting for image ID (*id* = root) as random factor (as one root per image were imaged, but five cells per root used). We then computed post hoc tests (*lsmeans* package

**Supplementary file 6.**
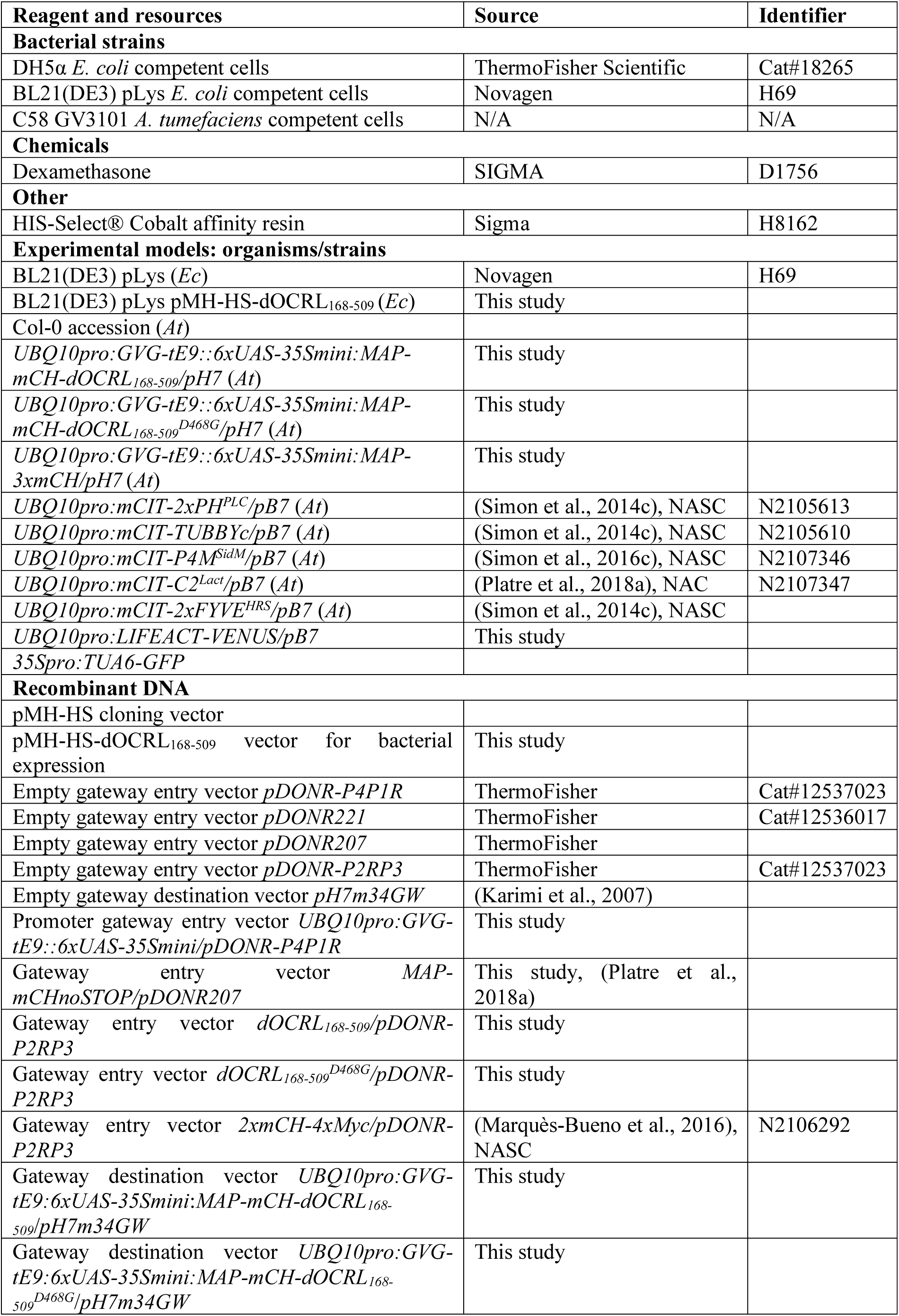
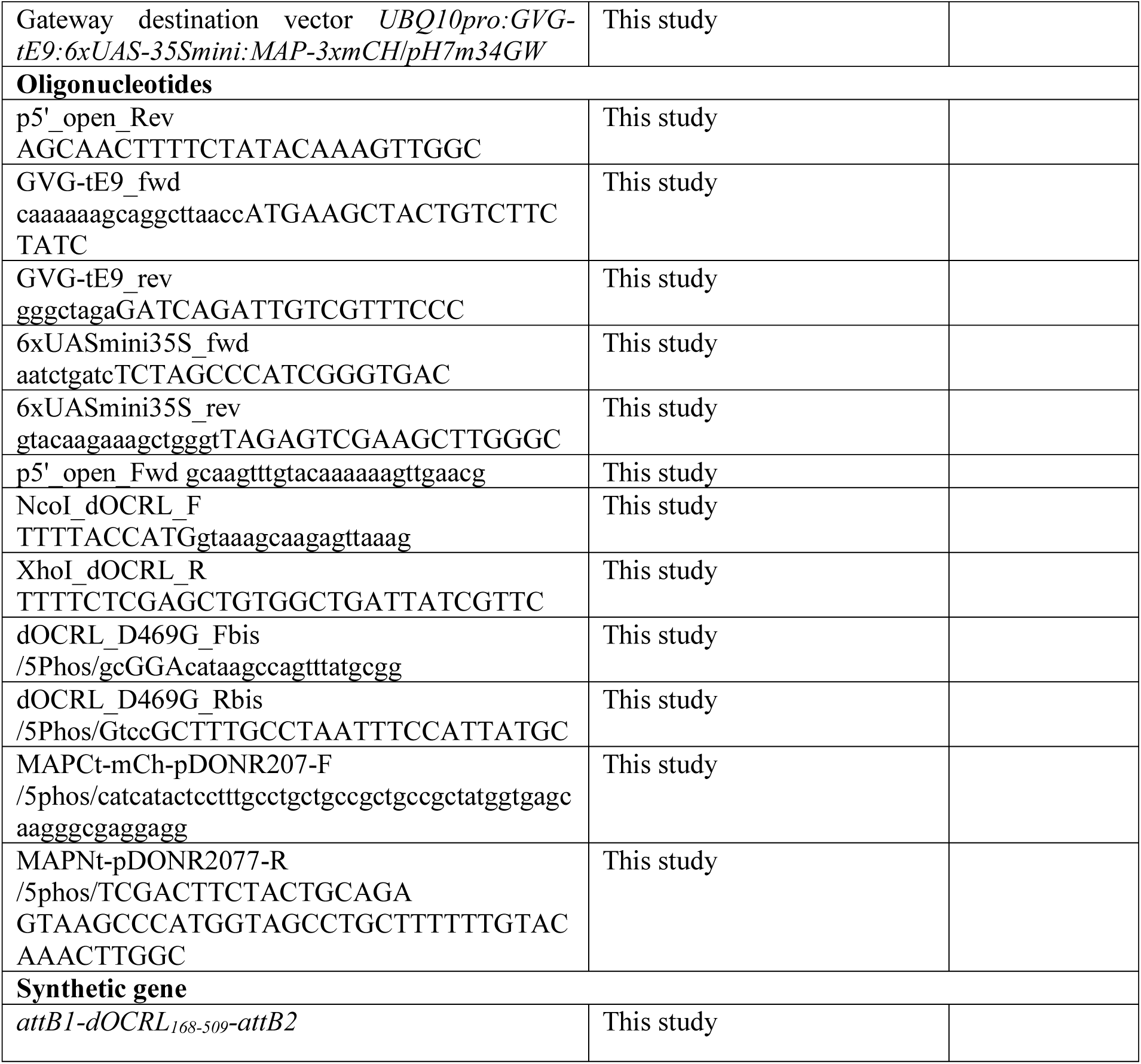
Reagent and resources.

